# CqFT1A and CqFT1B-1 are major flowering activators in quinoa

**DOI:** 10.64898/2026.07.07.736970

**Authors:** Takuya Ogata, Yasunari Fujita

## Abstract

Flowering time is a key determinant of crop adaptation, plant architecture, and breeding efficiency. Quinoa (*Chenopodium quinoa*), a climate-resilient allotetraploid orphan crop, exhibits extensive variation in flowering behavior, yet which of its multiple *FT* homologs drive flowering remains unclear. Here, we used virus-mediated overexpression (VOX) and virus-induced gene silencing (VIGS) in quinoa, together with heterologous expression in *Arabidopsis thaliana*, to dissect the flowering-promoting activity of quinoa *FLOWERING LOCUS T* (*FT*) homologs. Although multiple *CqFT* homologs were transcriptionally induced during the floral transition, their functional outputs were markedly unequal. *CqFT1A* and *CqFT1B-1* acted as the major florigenic activators: overexpression of either gene induced rapid and synchronized flowering, whereas the *CqFT1*-VIGS treatment delayed flowering. Although *CqFT2A* and *CqFT2B* showed weak flowering-promoting activity, their endogenous contribution remains unresolved because gene-specific silencing could not be achieved. *CqFT1B-2* had no detectable promotive effect under the conditions tested. Domain-swapping analyses revealed that replacing the C-terminal region of CqFT2A with that of CqFT1A partially restored flowering-promoting activity, indicating that C-terminal variation contributes to, but does not fully explain, functional divergence among *CqFT* homologs. In late-flowering highland lines, elevated FT input accelerated flowering, induced coordinated floral transition, and shortened the time to seed production. These findings identify *CqFT1A* and *CqFT1B-1* as the major florigenic activators in quinoa and demonstrate that transcriptional induction alone does not predict florigenic activity. More broadly, this work establishes a functional and methodological framework for resolving florigen activity in climate-resilient orphan crops of increasing agricultural importance.

## 1. Introduction

Flowering time integrates environmental signals with endogenous developmental programs and determines the transition from vegetative to reproductive growth (Andrés and Coupland, 2012). Because this transition governs reproductive success, crop duration, plant architecture, and adaptation to local environments, flowering time has been a major target of selection during domestication and remains a key trait for breeding regionally adapted and climate-resilient crops (Cockram et al., 2007; Eshed and Lippman, 2019; Jung and Muller, 2009). At the molecular level, flowering initiation is centrally regulated by the mobile florigen FLOWERING LOCUS T (FT), which transmits inductive signals from leaves to the shoot apical meristem (Kardailsky et al., 1999; Tamaki et al., 2007). In many crops, duplication and polyploidization have expanded *FT*-like gene families, creating paralogs with distinct, redundant, or even antagonistic functions (Higgins et al., 2010; Pin and Nilsson, 2012). Thus, determining which *FT* homologs provide the effective florigenic output is essential for understanding flowering-time control, particularly in polyploid crops.

Quinoa (*Chenopodium quinoa* Willd.) is an emerging orphan crop of global importance because of its high nutritional value and broad tolerance to abiotic stresses, including drought, salinity, frost, and low temperature (Jacobsen et al., 2005; Kobayashi and Fujita, 2024; Kobayashi et al., 2025; Mizuno et al., 2020; Vega-Gálvez et al., 2010; Yasui et al., 2016). Native to the Andean region, quinoa has diversified across contrasting environments, from lowland coastal areas to high-altitude plateaus, and exhibits pronounced variation in flowering behavior. Lowland accessions can flower rapidly under inductive short-day conditions, whereas several highland accessions display extreme late-flowering phenotypes that substantially prolong generation time (Bertero et al., 1999; Mizuno et al., 2020). This variation is central to quinoa adaptation but also constrains broader deployment of elite highland germplasm, including large-seeded “Real” types, outside their native environments (Bazile et al., 2016; Gomez-Pando et al., 2019; Schmöckel et al., 2017).

Genomic resources for quinoa have expanded rapidly. Draft and reference genome assemblies, followed by chromosome-scale genome assemblies and multi-genome comparative analyses, have provided increasingly detailed views of the allotetraploid quinoa genome and its flowering-time gene repertoire (Jarvis et al., 2017; Kobayashi et al., 2024; Patiranage et al., 2022; Rey et al., 2023; Rey et al., 2024; Yasui et al., 2016). These resources have revealed multiple *FT*-like homologs and enabled comparative analyses of flowering-related loci across diverse germplasm. However, genome sequence and expression information alone cannot determine which *FT* homologs function as effective floral activators *in planta*. In particular, it remains unclear whether flowering-time control in quinoa is distributed across multiple *FT* homologs or concentrated in a specific subset of *FT* homologs.

Functional dissection of flowering regulators in quinoa has been limited by its allotetraploid genome, genetic heterogeneity, and the lack of routine stable transformation or genome-editing systems. To address these limitations, we previously developed genotyped quinoa resources, including the genome-sequenced standard line Kd, to support reproducible genotype–phenotype analyses (Mizuno et al., 2020), and established apple latent spherical virus (ALSV)-based virus-induced gene silencing (VIGS) and virus-mediated overexpression (VOX) systems for quinoa, enabling transient loss- and gain-of-function analyses directly *in planta* (Ogata et al., 2021).

Here, we use ALSV-mediated VOX and VIGS, together with heterologous expression in *Arabidopsis thaliana*, to systematically evaluate the flowering-promoting activity of quinoa *FT* homologs and investigate the molecular basis of their functional divergence. We identify *CqFT1A* and *CqFT1B-1* as the major florigenic activators controlling flowering initiation in quinoa, whereas other *FT* homologs show weak or context-dependent activity. We further show that elevated *FT* input accelerates flowering across genetically and ecologically diverse quinoa lines, including strongly late-flowering highland accessions. These findings reveal a major contribution of the *CqFT1* homoeologous pair and provide a functional framework for modulating flowering time and generation turnover in this climate-resilient orphan crop.

## 2. Results

### 2.1 *CqFT* Homologs Are Differentially Expressed During the Floral Transition in Quinoa

To establish a foundation for functional dissection of flowering regulation in quinoa, we first characterized the genomic organization and developmental expression profiles of *FT* homologs. Genome annotation has identified five major *FT*-like genes in quinoa— *CqFT1A*, *CqFT1B-1*, *CqFT2A*, *CqFT2B*, and *CqFT1B-2* (Jarvis et al., 2017). Phylogenetic and sequence analyses indicate that *CqFT1A* and *CqFT1B-1*, as well as *CqFT2A* and *CqFT2B*, constitute homoeologous gene pairs derived from the A and B subgenomes, whereas *CqFT1B-2* is more distantly related (Figure 1a, Figure S1).

**FIGURE 1.**
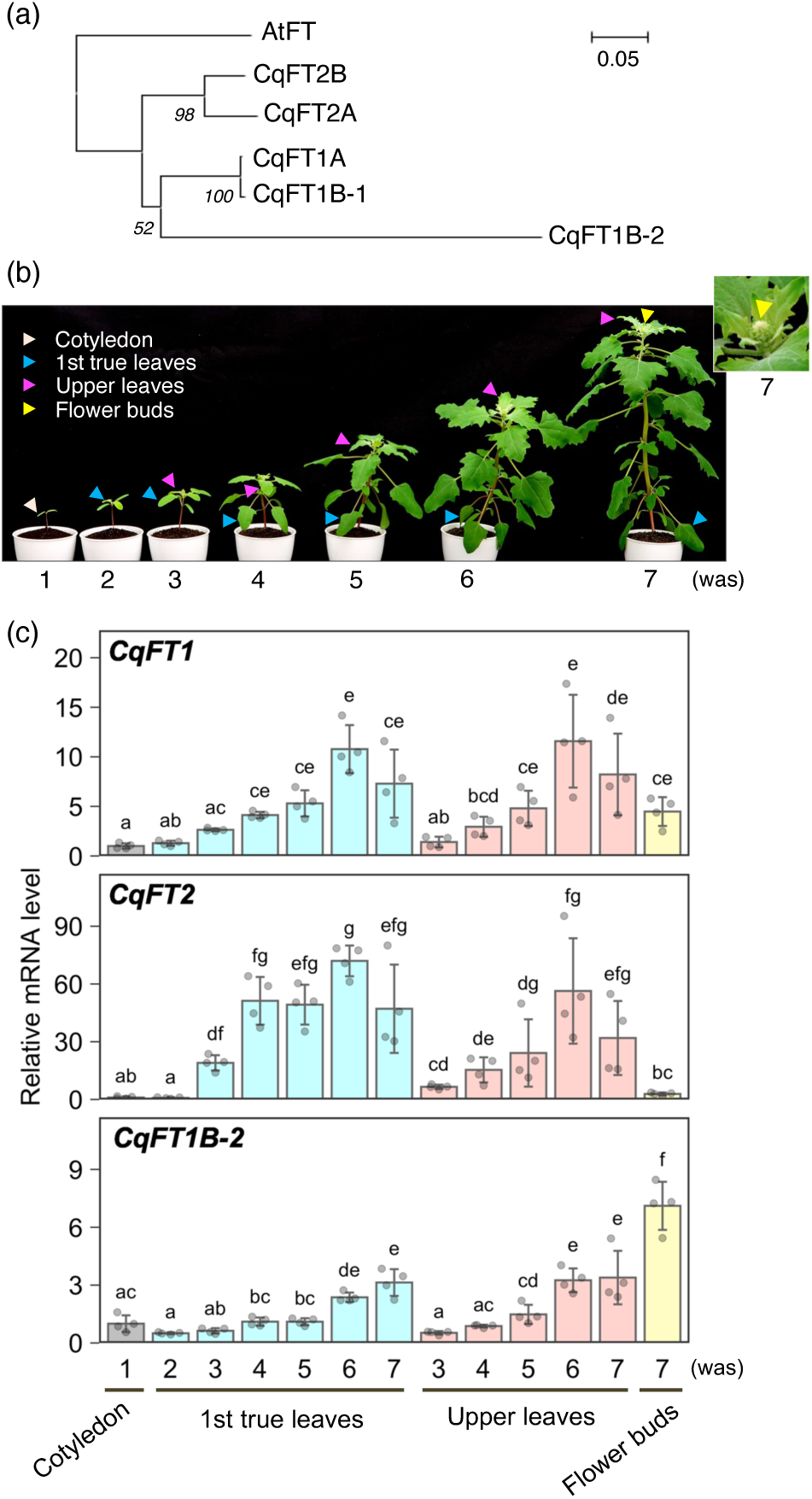
Developmental expression profiles of quinoa *FT* homologs during the floral transition. (a) Phylogenetic tree based on the protein sequences of quinoa CqFT homologs and *Arabidopsis thaliana* FT (AtFT). Sequences were aligned using ClustalW, and the tree was constructed using the neighbor-joining method. (b) Representative photographs of the lowland-type quinoa inbred line Kd, which was used for *CqFT* gene isolation and developmental expression analysis in this study. Plants were grown under a 12-h light/12-h dark photoperiod at 22°C during the light period and 16°C during the dark period. Cotyledons were sampled at 1 week after sowing (was), first true leaves at 2–7 was, upper leaves at 3–7 was, and flower buds at 7 was. Arrowheads indicate the tissues sampled at each time point. (c) RT-qPCR analysis of *CqFT1*, *CqFT2*, and *CqFT1B-2* transcript levels at the indicated growth stages. Primer pairs for *CqFT1* and *CqFT2* were designed within regions conserved between *CqFT1A*/*CqFT1B-1* and *CqFT2A*/*CqFT2B*, respectively, thereby detecting the combined transcript levels of each homoeologous gene pair. 18S *rRNA* was used for normalization. Data are presented as mean ± SD (*n* = 4 biological replicates). Different letters indicate significant differences (*p* < 0.01, Tukey’s HSD test on log-transformed data).

To examine how these *FT* homologs are regulated during the floral transition, we analyzed their expression dynamics in the leaves of the lowland quinoa line Kd (Yasui et al., 2016; Table S1) throughout vegetative growth and flowering induction. Full-length coding sequences of all five *CqFT* genes were isolated from the Kd plants, and their identities were confirmed by comparison with the QQ74 reference genome (Jarvis et al., 2017; Rey et al., 2023; Tables S1 and S2). Representative photographs of plants at each developmental stage are shown in Figure 1b; floral bud emergence occurred between 6 and 7 weeks after sowing. For each homoeologous gene pair (*CqFT1A*/*CqFT1B-1* and *CqFT2A*/*CqFT2B*), transcript abundance was quantified using primer sets that amplify both homoeologs (Table S3). Quantitative RT-PCR analyses revealed that transcripts of *CqFT1* (*CqFT1A*/*CqFT1B-1*) and *CqFT2* (*CqFT2A*/*CqFT2B*) were present at low levels during early vegetative growth but were progressively upregulated in leaves as plants approached flowering initiation (Figure 1c). For both *CqFT1* and *CqFT2*, transcript levels in leaves peaked at around week 6, with expression in flower buds at week 7 remaining below the week 6 leaf peak. In contrast, *CqFT1B-2* transcripts were progressively upregulated in leaves before flowering initiation, whereas expression in flower buds at week 7 exceeded that observed in leaves at weeks 6 and 7 (Figure 1c). Together, these results demonstrate that *CqFT* homologs in quinoa exhibit distinct temporal dynamics and tissue preferences of transcript accumulation during the floral transition, underscoring the necessity of functional analyses to resolve their respective contributions to flowering initiation.

### 2.2 *CqFT1A* and *CqFT1B-1* Accelerate Flowering in Quinoa

To determine whether individual quinoa *FT* homologs differ in their capacity to promote flowering, we conducted gain-of-function analyses using ALSV-mediated VOX in quinoa. Full-length coding sequences of *CqFT1A*, *CqFT1B-1*, *CqFT2A*, *CqFT2B*, and *CqFT1B-2* were independently expressed in young lowland quinoa plants (line J082) using the ALSV-based system (Table S1; Ogata et al., 2021). Because *Arabidopsis thaliana FT* (*AtFT*) is a well-established florigenic activator (Corbesier et al., 2007; Kardailsky et al., 1999), we first tested whether heterologous *AtFT* expression could induce flowering in quinoa. Plants inoculated with recombinant ALSV expressing *AtFT* exhibited a strong early-flowering phenotype, indicating that elevated *FT* input is sufficient to promote floral transition in quinoa (Figure S2a–c). We therefore used *AtFT*-VOX as a positive control for flowering induction in subsequent VOX assays.

Plants inoculated with a green fluorescent protein (GFP)-expressing virus or an empty vector served as negative controls. Quinoa plants overexpressing *CqFT1A* or *CqFT1B-1* exhibited markedly accelerated flowering compared with control plants, although the acceleration was less pronounced than that observed in *AtFT*-overexpressing plants (Figure 2a–c). In contrast, overexpression of *CqFT2A*, *CqFT2B*, or *CqFT1B-2* did not significantly alter flowering time relative to controls under the conditions tested (Figure 2b,c), despite confirmed transgene expression (Figure 2d–f). Notably, in plants overexpressing *AtFT*, *CqFT1A*, or *CqFT1B-1*, floral transition occurred synchronously across the plant: both primary and axillary shoot apices developed flower buds within the same observational time window, in contrast to the sequential or apex-restricted pattern seen in controls (Figure S2d–f). In control and non-accelerated plants, floral bud formation was confined to the primary shoot apex, while the majority of axillary meristems remained vegetative throughout the observation period. Collectively, these results demonstrate that among the quinoa *FT* homologs tested, only *CqFT1A* and *CqFT1B-1* function as effective promoters of flowering in quinoa, revealing functional divergence among transcriptionally induced *CqFT* homologs.

**FIGURE 2.**
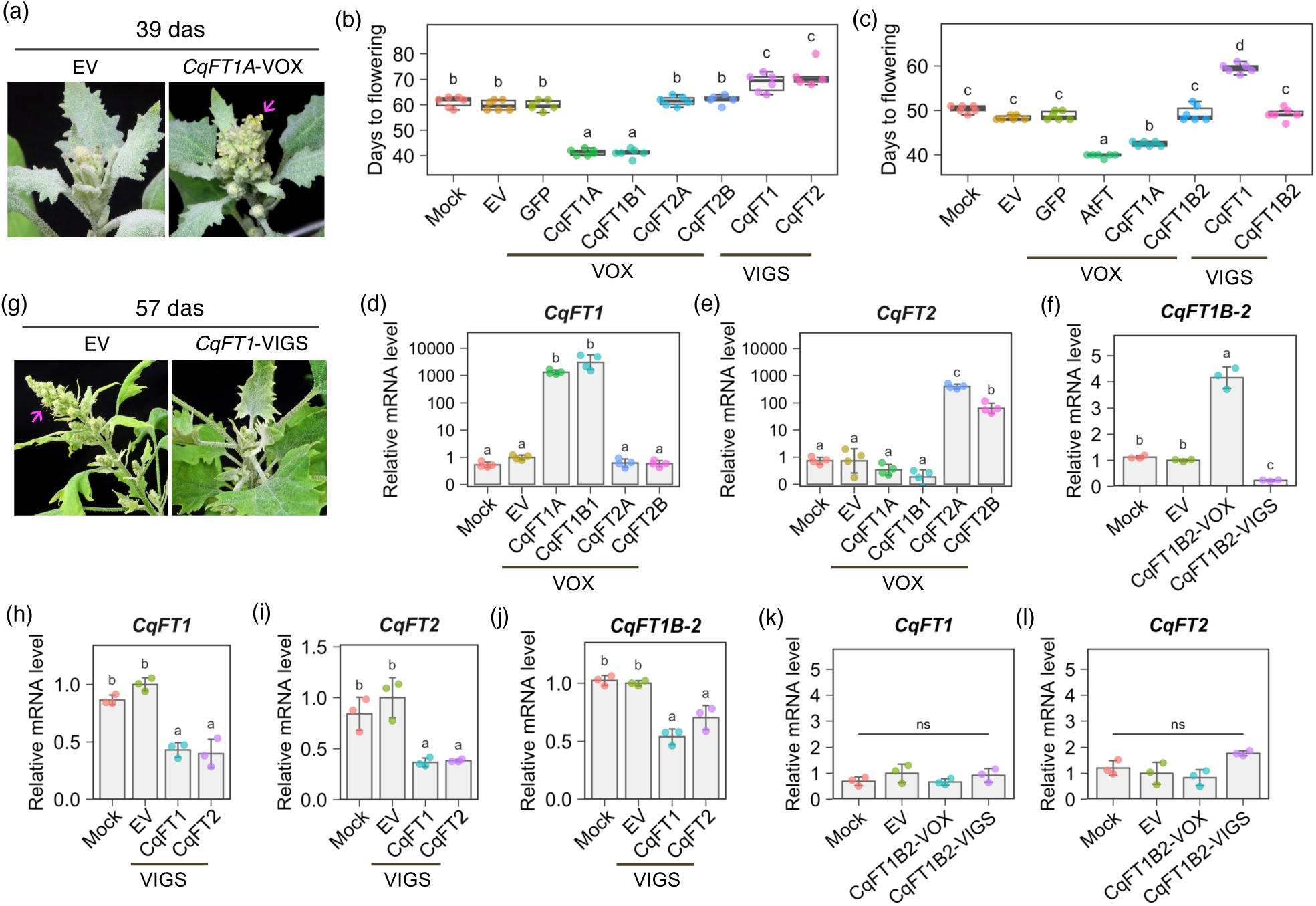
ALSV-mediated VOX and VIGS of quinoa *FT* homologs modulate flowering time in quinoa. (a) Representative shoot apices of quinoa J082 plants inoculated with recombinant ALSV inocula corresponding to the empty vector (EV) or *CqFT1A*-VOX at 39 days after sowing (das). Arrows indicate opening flower buds. (b, c) Days to flowering in plants subjected to the indicated VOX or VIGS treatments. Days to flowering were recorded as the number of days after sowing until at least one flower bud opened. Buffer-only mock inoculation, EV, and GFP-VOX were used as controls. Data are presented as mean ± SD (*n* = 6 plants). Different letters indicate significant differences (*p* < 0.01, Tukey’s HSD test). (d–f) RT-qPCR analysis of *CqFT1* (d), *CqFT2* (e), and *CqFT1B-2* (f) transcript levels in plants subjected to the indicated VOX or VIGS treatments. Data are presented as mean ± SD (*n* = 4 biological replicates). Different letters indicate significant differences (*p* < 0.01, Tukey’s HSD test on log-transformed data). (g) Representative shoot apices of plants inoculated with recombinant ALSV inocula corresponding to EV or *CqFT1*-VIGS at 57 das. Arrows indicate opening flower buds. (h–j) RT-qPCR analysis of *CqFT1* (h), *CqFT2* (i), and *CqFT1B-2* (j) transcript levels in plants subjected to *CqFT1*- or *CqFT2*-VIGS. Data are presented as mean ± SD (*n* = 3 biological replicates). Different letters indicate significant differences (*p* < 0.05, Tukey’s HSD test). (k, l) RT-qPCR analysis of *CqFT1* (k) and *CqFT2* (l) transcript levels in plants subjected to *CqFT1B-2*-VOX or *CqFT1B-2*-VIGS. Data are presented as mean ± SD (*n* = 3 biological replicates). Different letters indicate significant differences (*p* < 0.01, Tukey’s HSD test on log-transformed data); ns indicates not significant. Upper leaf samples for RT-qPCR were collected at 20 days after ALSV inoculation. *CqUBQ10* was used for normalization. For *CqFT1* and *CqFT2*, primer pairs detected the combined transcript levels of *CqFT1A*/*CqFT1B-1* and *CqFT2A*/*CqFT2B*, respectively.

### 2.3 *FT*-Targeting VIGS Delays Flowering in Quinoa

To complement the gain-of-function analyses and assess the contribution of endogenous *CqFT* activity to flowering time in quinoa, we next performed gene-silencing analyses using VIGS. Because of the high sequence similarity between homoeologous gene pairs, *CqFT1A* and *CqFT1B-1*, as well as *CqFT2A* and *CqFT2B*, were simultaneously targeted using VIGS constructs designed to silence each homoeologous pair (Figure S3). For *CqFT1B-2*, a gene-specific region was selected to enable independent silencing (Figure S3). Plants inoculated with the *CqFT1*-targeting VIGS inoculum showed a significant delay in flowering compared with control plants (Figure 2b,c,g). *CqFT2*-VIGS also delayed flowering (Figure 2b); however, because this treatment concomitantly reduced *CqFT1* transcript levels (Figure 2h,i), the delayed flowering observed in plants subjected to *CqFT2*-VIGS cannot be unambiguously attributed to the loss of *CqFT2* function alone but may partly reflect the concomitant reduction of *CqFT1*. Although the VOX results indicate that *CqFT1A* and *CqFT1B-1* act as key accelerators of flowering in quinoa (Figure 2a,b,c), a possible contribution of endogenous *CqFT2A*/*CqFT2B* to flowering initiation cannot be excluded because independent silencing could not be achieved. To improve the target specificity of *CqFT1* and *CqFT2* silencing, we attempted VIGS using sequences derived from the 5′ or 3′ untranslated regions or concatenated untranslated region (UTR) fragments. These approaches, however, did not achieve effective or reproducible suppression and failed to induce measurable changes in flowering time (Figure S4). In contrast, although *CqFT1B-2*-VIGS specifically suppressed *CqFT1B-2* expression (Figure 2f,k,l), it had no detectable effect on the timing of flowering (Figure 2b,c).

### 2.4 Quinoa FT Homologs Exhibit Differential Flowering-Promoting Activity in Arabidopsis

To evaluate the flowering-promoting activity of quinoa *FT* homologs in an independent stable heterologous expression system, we generated transgenic *Arabidopsis* lines expressing individual *CqFT* coding sequences under the control of the cauliflower mosaic virus 35S (CaMV 35S) promoter. Non-transgenic wild-type Col-0 plants and transgenic lines expressing GFP under the CaMV 35S promoter (35S-GFP) were used as negative controls, whereas transgenic lines expressing *AtFT* under the CaMV 35S promoter (35S-*AtFT*) served as positive controls. In the T1 generation, a large proportion of independent transformants expressing 35S-*CqFT1A* or 35S-*CqFT1B-1* displayed a striking early-flowering phenotype on selection media, with many plants initiating bolting at the cotyledon stage before the emergence of true leaves (Figure 3a). In contrast, none of the T1 transformants expressing 35S-*CqFT2A*, 35S-*CqFT2B*, or 35S-*CqFT1B-2* exhibited conspicuous early flowering under the same conditions (Figure 3a).

**FIGURE 3.**
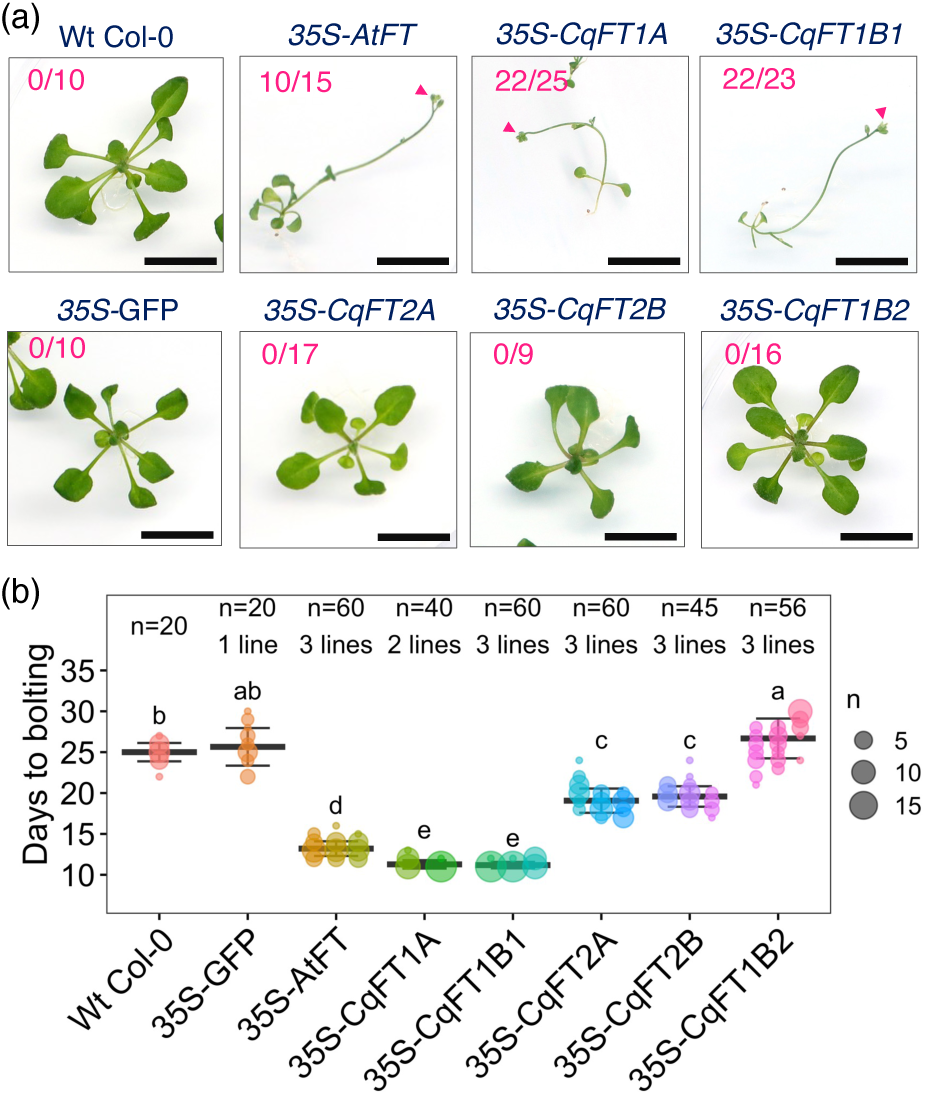
Quinoa *FT* homologs exhibit differential flowering-promoting activity in *Arabidopsis*. *Arabidopsis thaliana* Col-0 was transformed with CaMV 35S promoter-driven *FT* genes. Wild-type (Wt) Col-0 and T3 homozygous GFP-overexpressing lines (35S-GFP) were grown as controls. (a) Photographs were taken 19 days after germination day 0, defined as the day of transfer to 22°C. The number of independent T1 transformants that had initiated flowering relative to the total number of T1 transformants generated is shown. For Wt Col-0 and 35S-GFP controls, 10 plants were analyzed, and none had initiated flowering at this time point. Scale bar = 1 cm. (b) Days to bolting from germination day 0 were recorded for each line. Two or three independent T2 lines were analyzed per construct. Data are presented as mean ± SD of all plants analyzed, and individual lines are shown in different colors. The number of lines and total plants (n) are indicated at the top. Different letters indicate significant differences (*p* < 0.001, Dunnett’s T3 test).

To quantitatively assess flowering time, two to three independent T2 lines were selected for each construct. Transgene expression was confirmed by RT-PCR (Figure S5a). Measurements of days to bolting, days to flowering, and total leaf number at flowering consistently showed that 35S-*CqFT1A* and 35S-*CqFT1B-1* lines flowered significantly earlier than all controls, including 35S-*AtFT* lines, bolting at 11–12 days after sowing (Figure 3b, Figure S5b,c). By contrast, 35S-*CqFT2A* and 35S-*CqFT2B* lines exhibited only modest flowering acceleration, and their flowering times remained substantially later than those of 35S-*AtFT*, 35S-*CqFT1A*, or 35S-*CqFT1B-1* lines (Figure 3b, Figure S5b,c). Notably, the modest but significant early-flowering activity of *CqFT2A* and *CqFT2B* in *Arabidopsis* contrasts with their lack of detectable effect in quinoa VOX experiments (Figure 2b,c), indicating that these homologs retain weak flowering-promoting capacity that was not readily detected in quinoa VOX assays. Expression of 35S-*CqFT1B-2* did not accelerate flowering and, in some lines, showed a slight delay relative to controls (Figure 3b, Figure S5b,c). Taken together, these results demonstrate that quinoa *FT* homologs differ markedly in flowering-promoting activity in *Arabidopsis*: *CqFT1A* and *CqFT1B-1* exhibit strong activity, *CqFT2A* and *CqFT2B* display weak activity, and *CqFT1B-2* lacks detectable promoting capacity.

### 2.5 Altered *FT* Expression Modifies Vegetative Growth and Final Plant Height

Beyond its pronounced effects on flowering time, manipulation of *FT* expression markedly altered vegetative growth dynamics in quinoa. Plants in which flowering was strongly accelerated exhibited enhanced vertical growth prior to the floral transition, resulting in greater plant height at the onset of flowering compared with control plants (Figure 4a). This transient increase in height was primarily attributable to an increased number of nodes rather than elongation of internodes (Figure S6a,b), indicating that flowering acceleration modifies the timing of developmental phase progression without promoting generalized stem elongation. Following the initiation of flowering, however, vegetative growth ceased earlier in early-flowering plants, leading to a reversal of height differences at later developmental stages. Consequently, final plant height at maturity was reduced in early-flowering plants relative to controls (Figure 4b,c; Figure S6c).

**FIGURE 4.**
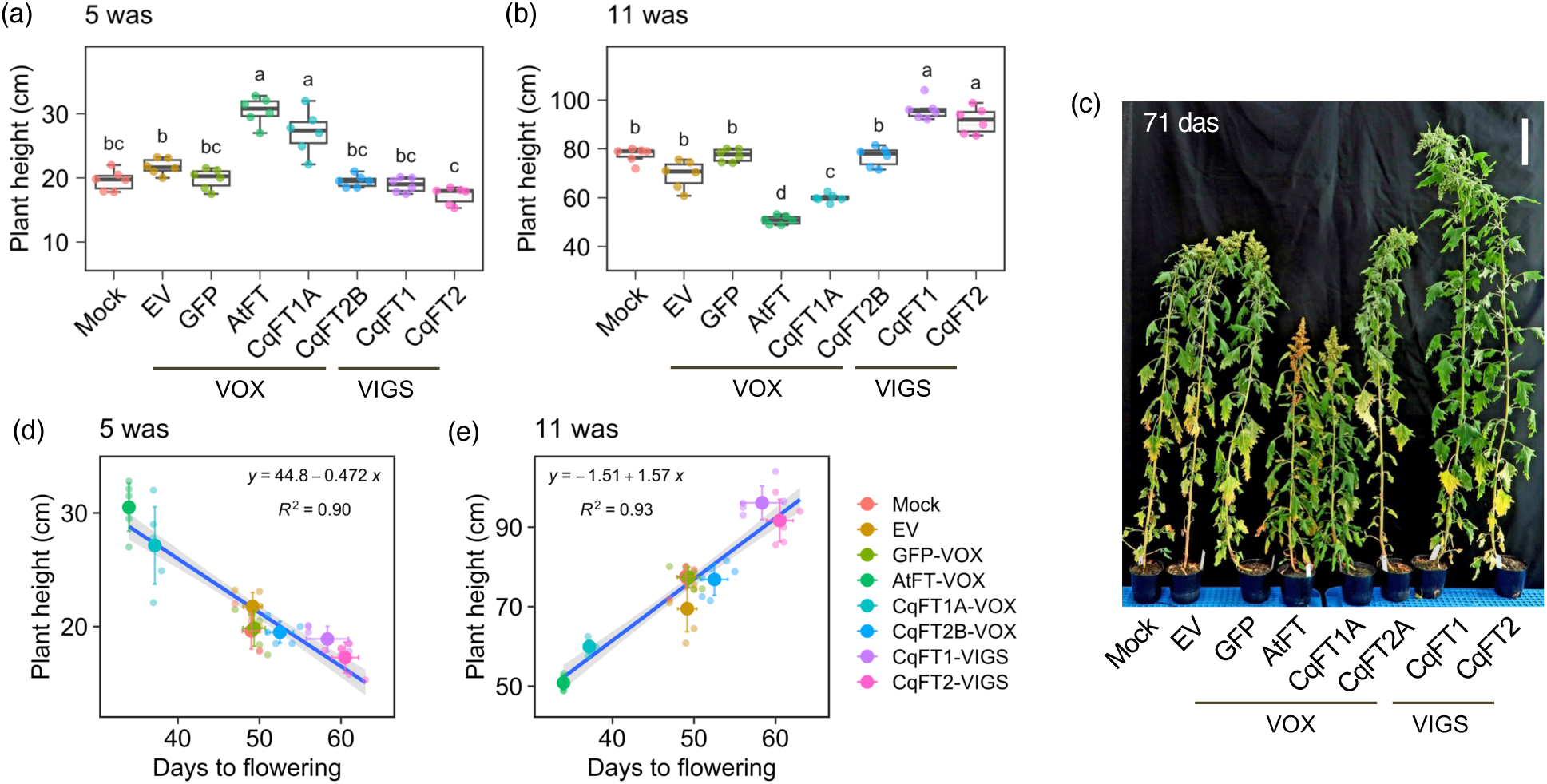
Altered *FT* expression modifies flowering time and plant height in quinoa. (a, b) Plant height of quinoa J082 plants subjected to ALSV-mediated VOX or VIGS of *FT* genes. Plant height was measured at 5 weeks after sowing (was) (a) and 11 was (b). Buffer-only mock inoculation, EV, and GFP-VOX were used as controls. Data are presented as mean ± SD (*n* = 6 plants). Different letters indicate significant differences (*p* < 0.01, Tukey’s HSD test). (c) Representative photograph of plants subjected to the indicated VOX or VIGS treatments. The photograph was taken at 71 days after sowing (das), one week after flowering initiation in *CqFT1*-VIGS plants. Scale bar = 10 cm. (d, e) Relationship between flowering time and plant height at 5 was (d) and 11 was (e). Mean days to flowering for each treatment were plotted against mean plant height at the indicated time point. Data are presented as mean ± SD (*n* = 6 plants). Linear regression models and coefficients of determination (R²) are shown.

Conversely, suppression of endogenous flowering induction prolonged the vegetative phase and delayed floral transition. These plants continued vertical growth for a longer period and ultimately attained greater final height than control plants (Figure 4b,c). As a result, the relationship between plant height and flowering time shifted dynamically during development: early flowering was associated with transient height increases but reduced final stature, whereas delayed flowering resulted in prolonged vegetative growth and increased final height (Figure 4d,e). Together, these results indicate that *FT*-mediated changes in flowering time are closely associated with changes in vegetative growth duration and final plant height, linking floral transition timing to whole-plant developmental trajectories in quinoa.

### 2.6 C-terminal Sequence Variation Contributes to Functional Divergence among Quinoa FT Homologs

Although multiple *CqFT* homologs are transcriptionally induced during the floral transition with distinct temporal dynamics (Figure 1), gain- and loss-of-function analyses revealed substantial differences in their flowering-promoting capacities (Figures 2, 3). To elucidate the molecular basis of this functional divergence, we compared amino acid sequences of CqFT proteins exhibiting contrasting flowering activities. Alignment of full-length FT proteins revealed variation in both sequence length and composition among quinoa FT homologs (Figure 5a). Relative to the strong flowering promoters CqFT1A and CqFT1B-1, CqFT2A and CqFT2B lacked four amino acids at the N-terminus, whereas CqFT1B-2 possessed markedly extended N- and C-terminal regions. Among all amino acid substitutions, twelve residues were uniquely shared by the weak or non-promoting FT homologs (CqFT2A/2B and CqFT1B-2) but absent from the strong flowering promoters AtFT, CqFT1A, and CqFT1B-1. Several of these substitutions mapped to regions previously implicated in FT function, including the conserved segment C and the unstructured C-terminal tail (Ahn et al., 2006; Gao et al., 2025; Ho and Weigel, 2014; Wang et al., 2017).

**FIGURE 5.**
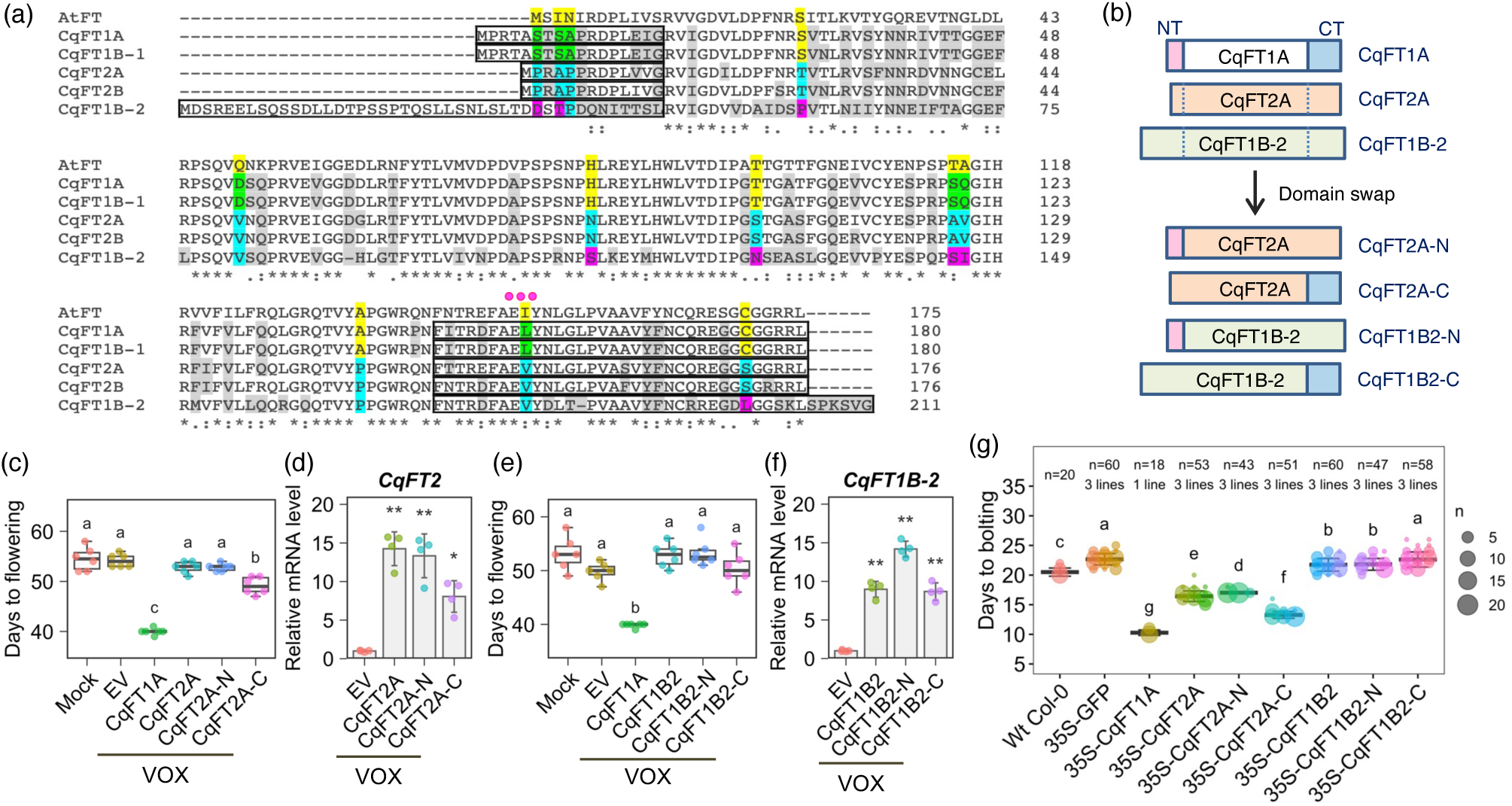
C-terminal sequence variation contributes to differential flowering-promoting activity among quinoa FT homologs. (a) Multiple sequence alignment of AtFT and CqFT proteins generated using ClustalW. Amino acid residues shared by CqFT2A/CqFT2B and CqFT1B-2, but not by CqFT1A/CqFT1B-1 or AtFT, are highlighted in cyan or magenta. Swapped N-terminal (NT) and C-terminal (CT) regions are indicated. The conserved LYN/IYN triplets in segment C of exon 4 (Ahn et al., 2006) are marked with magenta dots. (b) Schematic diagram of domain-swapped CqFT constructs. The NT or CT regions of CqFT2A and CqFT1B-2 were replaced with the corresponding regions of CqFT1A. (c, e) Days to flowering in quinoa plants subjected to VOX of native or domain-swapped CqFT2A (c) and CqFT1B-2 (e). Buffer-only mock inoculation and EV were used as controls. Data are presented as mean ± SD (*n* = 6 plants). Different letters indicate significant differences (*p* < 0.01, Tukey’s HSD test). (d, f) RT-qPCR analysis of transgene expression in quinoa plants subjected to VOX of native or domain-swapped CqFT2A (d) and CqFT1B-2 (f), using primer pairs for *CqFT2* and *CqFT1B-2*, respectively. Upper leaf samples were collected at 17 days after ALSV inoculation. *CqUBQ10* was used for normalization. Data are presented as mean ± SD (*n* = 4 biological replicates). Statistical significance was determined using Welch’s *t*-test with Bonferroni correction on log-transformed data (\**p* < 0.05, \*\**p* < 0.01). (g) Days to bolting in transgenic *Arabidopsis* lines expressing native or domain-swapped CqFT constructs under the CaMV 35S promoter. Days to bolting from germination day 0 were recorded for each line. Three independent T2 lines were analyzed per construct, except for the 35S-*CqFT1A* line used as a control. Wt Col-0 and 35S-GFP were grown as controls. Data are presented as mean ± SD of all plants analyzed, and individual lines are shown in different colors. Different letters indicate significant differences (*p* < 0.001, Dunnett’s T3 test).

To directly test whether terminal regions contribute to FT functional divergence, we generated domain-swapping constructs in which either the N- or C-terminal regions of CqFT2A or CqFT1B-2 were replaced with the corresponding regions of CqFT1A (Figure 5b). When expressed via VOX in quinoa, substitution of the N-terminal region of CqFT2A had no detectable effect on flowering time relative to the native protein (Figure 5c,d). In contrast, substitution of the C-terminal region conferred a modest but statistically significant increase in flowering-promoting activity, advancing flowering by approximately 4 days relative to wild-type CqFT2A, although flowering remained substantially later than in CqFT1A-overexpressing plants (Figure 5c,d). Domain swapping of either terminal region of CqFT1B-2 did not confer detectable flowering-promoting activity (Figure 5e,f). To validate these findings in an independent system and exclude potential effects of viral co-suppression, we generated transgenic *Arabidopsis* lines expressing the domain-swapped constructs under the CaMV35S promoter. Consistent with the quinoa VOX results, plants expressing the C-terminal–swapped version of CqFT2A flowered earlier than those expressing native CqFT2A, yet later than lines expressing CqFT1A (Figure 5g, Figure S7). In contrast, *Arabidopsis* lines expressing domain-swapped versions of CqFT1B-2 showed little or no effect on flowering time (Figure 5g, Figure S7).

Within the 34 amino acids comprising the swapped C-terminal region, five residues distinguished CqFT2A from CqFT1A. Notably, two substitutions (I/L151V and C171S/L) consistently segregated strong flowering promoters from weak or inactive FT homologs (Figure 5a). Computational structural prediction using ColabFold program suggested that the conserved segment C adopts a stable fold, whereas the C-terminal region exhibits low confidence scores, consistent with intrinsic structural flexibility (Figure S8). The C171 residue is located within this low-confidence region, suggesting that the C-terminal tail may be structurally flexible. Collectively, these results demonstrate that C-terminal sequence variation contributes to functional divergence among quinoa FT homologs, although its contribution varies among paralogs and additional determinants are likely involved.

### 2.7 Altered *FT* Expression Modifies Flowering Time across Diverse Quinoa Genetic Backgrounds

To determine whether altered *FT* expression modifies flowering time across genetically diverse quinoa lines, we examined the effects of increased and reduced *FT* expression in highland-type quinoa lines that exhibit pronounced late-flowering phenotypes. The highland lines J067, J131, and J134 were selected for analysis because they exhibit markedly delayed bolting and flowering compared with lowland lines under short-day conditions (12 h light/12 h dark) (Mizuno et al., 2020). Consistent with previous reports, these highland lines initiated bolting substantially later than the lowland lines Kd and J082, with J134 displaying the most extreme delay, bolting approximately 30 days later than J082 (Figure 6a,b, Figure S9a,b). Analysis of endogenous *FT* expression revealed that *CqFT1* transcripts were induced at the time of bolting initiation in all lines, but expression levels in the highland lines were markedly lower than in the lowland lines (Figure 6c). *CqFT2* expression exhibited a similar induction pattern in J067, whereas its upregulation was weak or inconsistent in the strongly late-flowering lines J131 and J134 (Figure 6d). In contrast, *CqFT1B-2* transcripts were induced in all lines, but expression levels in J067 and J131 remained significantly lower than those observed in J134 and in the lowland lines (Figure 6e). Notably, *CqFT1B-2* expression in J134 was markedly higher than in the other highland lines, suggesting a potentially distinct regulatory context in this most strongly late-flowering accession, although the functional significance of this elevated expression remains to be determined.

**FIGURE 6.**
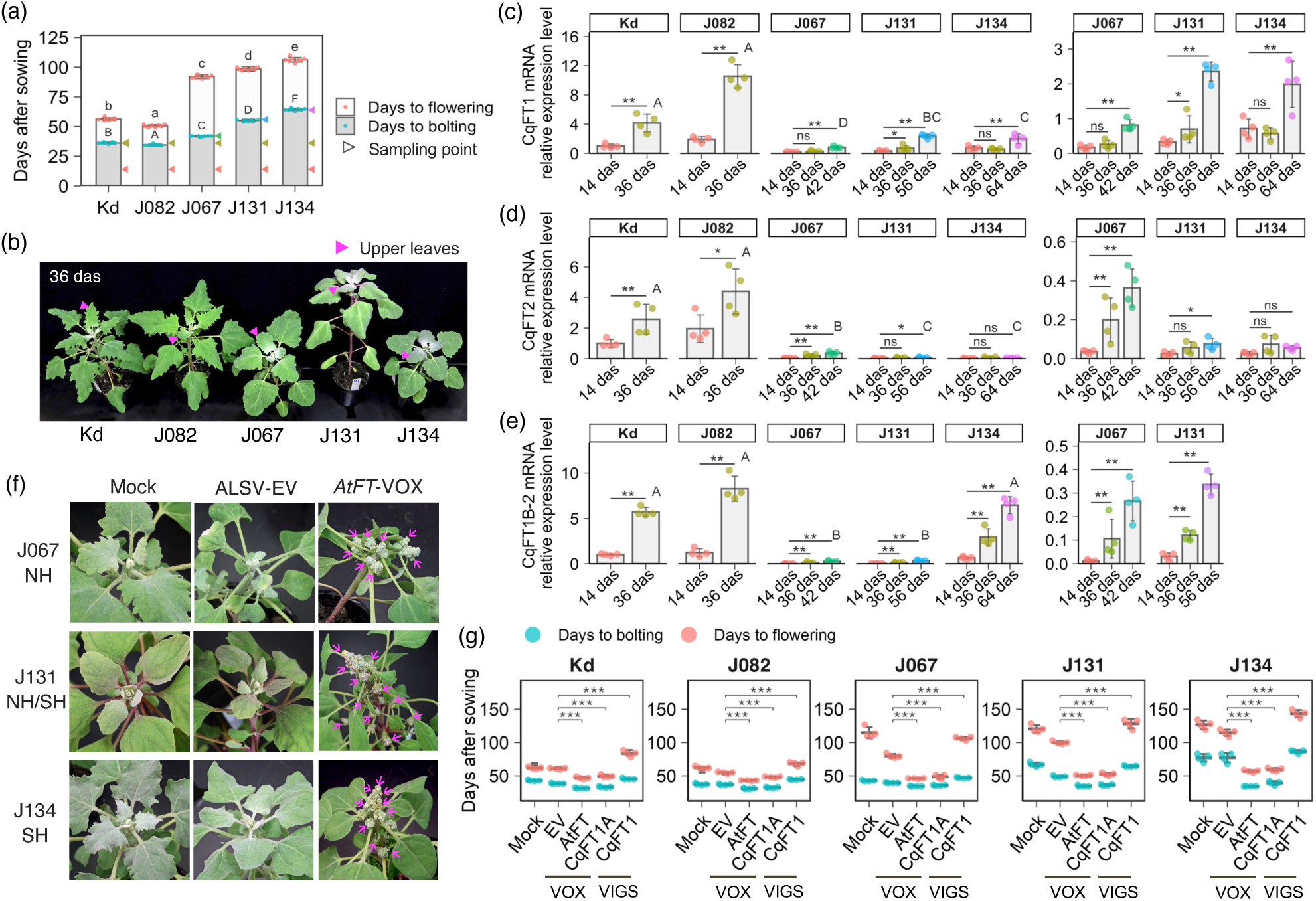
Altered *FT* expression modifies flowering time across diverse quinoa genetic backgrounds. (a) Comparison of bolting and flowering time in quinoa inbred lines Kd, J082, J067, J131, and J134 grown under short-day conditions. Plants were grown under a 12-h light/12-h dark photoperiod at 22°C during the light period and 16°C during the dark period. Data are presented as mean ± SD (*n* = 7 plants). Different letters indicate significant differences (*p* < 0.01, Tukey’s HSD test). Upper leaf samples were collected on the days indicated by arrowheads. (b) Representative photographs at 36 days after sowing (das), when the lowland lines Kd and J082 began bolting. Arrowheads indicate sampled upper leaves. Photographs from other sampling points are shown in Figure S9a,b. (c–e) RT-qPCR analysis of endogenous *CqFT* transcript levels in the five inbred lines. Leaf samples were collected at the time points shown in (a). Transcript levels were normalized to *CqUBQ10* and expressed relative to Kd at 14 das for *CqFT1* (c), *CqFT2* (d), and *CqFT1B-2* (e). Data are presented as mean ± SD (*n* = 4 biological replicates). Asterisks indicate significant differences from the Kd 14-das reference as determined by Dunnett’s test (\**p* < 0.05, \*\**p* < 0.01; ns, not significant). Different letters indicate significant differences among lines at 36 das for Kd and J082, 42 das for J067, 56 das for J131, and 64 das for J134, as determined by Tukey’s HSD test (*p* < 0.01). Data were log-transformed for statistical analyses. Enlarged y-axis scales for J067, J131, and J134 are shown on the right. For *CqFT1* and *CqFT2*, primer pairs detected the combined transcript levels of *CqFT1A*/*CqFT1B-1* and *CqFT2A*/*CqFT2B*, respectively. (f) Inoculation with recombinant ALSV inocula corresponding to *AtFT*-VOX or *CqFT1A*-VOX accelerates flowering in late-flowering highland quinoa lines. Inoculated plants were grown under short-day conditions with a 12-h light period at 22°C and a 12-h dark period at 20°C. Representative photographs at 50 das are shown. Arrows indicate opening flower buds. Buffer-only mock inoculation and EV were used as controls. (g) Bolting and flowering time in late-flowering highland lines inoculated with recombinant ALSV inocula corresponding to EV, *AtFT*-VOX, *CqFT1A*-VOX, or *CqFT1*-VIGS. Data are presented as mean ± SD (*n* = 4 plants). Statistical significance was determined using Dunnett’s test (\*\*\**p* < 0.001 versus EV).

To directly test the effects of altered *FT* expression on flowering time in these late-flowering backgrounds, ALSV-based VOX and VIGS constructs were inoculated into the highland lines at 14 days after sowing. In all three highland backgrounds, plants expressing *AtFT* or *CqFT1A* exhibited a pronounced acceleration of bolting and flowering relative to empty-vector controls, with flowering initiated 20–30 days earlier and accompanied by synchronous floral transition across both primary and axillary meristems (Figure 6f,g, Figure S9c–h). Conversely, plants subjected to *CqFT1*-VIGS delayed flowering in all three highland backgrounds, although the magnitude of delay varied among lines (Figure 6g, Figure S9f–h), consistent with a contribution of endogenous *CqFT* activity to flowering-time regulation. Plants subjected to *AtFT*- or *CqFT1A*-mediated flowering acceleration completed reproductive development and produced viable seeds within a shortened timeframe across all genetic backgrounds tested (Figure S10). Collectively, these findings demonstrate that altered *FT* expression modifies flowering time across genetically and ecologically distinct quinoa lines. Despite substantial variation in endogenous *FT* expression levels and flowering behavior, increased *FT* expression consistently accelerated flowering, whereas reduced *FT* expression delayed flowering in all tested highland backgrounds. These findings suggest that manipulation of *FT* expression may provide a useful approach for modifying flowering behavior across diverse quinoa genetic backgrounds.

## 3. Discussion

Here, we show that *CqFT1A* and *CqFT1B-1* function as the major florigenic activators in quinoa (Figures 2 and 3; Figures S2 and S5), despite the transcriptional activation of multiple *FT* homologs before flowering (Figure 1c). This conclusion could not have been reached from genome annotation or expression profiles alone (Golicz et al., 2020; Jarvis et al., 2017; Patiranage et al., 2021). Rather, it required direct functional testing using complementary gain- and loss-of-function approaches in quinoa, together with heterologous expression in *Arabidopsis*. Overexpression of either *CqFT1A* or *CqFT1B-1* was sufficient to induce rapid and synchronized flowering, whereas inoculation with the *CqFT1*-targeting VIGS construct delayed flowering, supporting the *CqFT1* homoeologous pair as a determinant of endogenous flowering time in quinoa. Consistent with the conserved role of FT-family florigens as integrators of flowering pathways (Takagi et al., 2023; Turck et al., 2008), similar specialization among FT paralogs has been reported in soybean, wheat, and rice, where only a subset of FT family members functions as major floral promoters (Halliwell et al., 2016; Komiya et al., 2008; Kong et al., 2010; Liu et al., 2018; Lv et al., 2014; Tamaki et al., 2007).

A notable finding of this study is that transcriptional induction was a poor predictor of florigenic activity among quinoa *FT* homologs. Although *CqFT1*, *CqFT2*, and *CqFT1B-2* were all induced during the floral transition, only *CqFT1A*/*CqFT1B-1* exhibited strong flowering-promoting activity (Figures 1–3; Figures S2 and S5). *CqFT2A*/*CqFT2B* exhibited weak flowering-promoting activity, whereas *CqFT1B-2* exhibited little or no detectable activity in our assays. The endogenous contribution of *CqFT2A*/*CqFT2B* remains unresolved because independent loss-of-function analysis could not be achieved. Nevertheless, the lack of detectable flowering acceleration upon *CqFT2A*/*CqFT2B* overexpression in quinoa, together with their modest activity in *Arabidopsis*, suggests that these homologs do not possess strong stand-alone flowering-promoting activity under the conditions tested. Thus, the differential activities of quinoa *FT* homologs point to functional differences that arise downstream of transcriptional induction.

Domain-swapping analyses indicate that protein-level variation contributes to the unequal florigenic activity of quinoa FT homologs (Figure 5, Figure S7). Replacing the C-terminal region of the weak activator CqFT2A with the corresponding region of CqFT1A modestly but significantly increased flowering-promoting activity, whereas N-terminal substitution had no detectable effect. However, the incomplete recovery of CqFT1A-level activity indicates that C-terminal variation explains only part of the functional divergence between these paralogs. Moreover, substitution of either terminal region failed to confer detectable flowering-promoting activity on CqFT1B-2, suggesting that additional determinants outside the swapped regions are involved, potentially including alterations in the conserved IYN motif implicated in FT/TFL1 activity switching (Ahn et al., 2006). These results are consistent with previous studies showing that short surface-exposed motifs and C-terminal residues in FT/TFL1 proteins modulate interactions with FD or 14-3-3 proteins and alter floral regulatory activity (Gao et al., 2025; Ho and Weigel, 2014; Wang et al., 2017; Wickland and Hanzawa, 2015). Thus, functional divergence among quinoa FT homologs appears to arise, at least in part, from quantitative tuning of florigenic activity at the protein level.

The highland lines examined here showed reduced or delayed endogenous induction of *CqFT1*, yet remained responsive to elevated *FT* input (Figure 6, Figure S9). Viral overexpression of *AtFT* or *CqFT1A* accelerated flowering in all tested highland backgrounds and induced a coordinated floral transition across primary and axillary meristems. Conversely, plants subjected to *CqFT1*-VIGS showed delayed flowering across these backgrounds, consistent with a contribution of endogenous *CqFT* activity to flowering-time regulation. These findings suggest that late flowering behavior in the lines examined is associated, at least in part, with differences in *CqFT1* expression regulation rather than a loss of downstream competence to respond to florigen. Such retained *FT* responsiveness provides a basis for modulating flowering time across genetically and ecologically distinct quinoa materials.

Manipulating *FT* activity also revealed a close coupling between flowering time and whole-plant developmental trajectories. Accelerated flowering was associated with earlier cessation of vegetative growth and reduced final stature, whereas delayed flowering prolonged vegetative growth and increased plant height (Figure 4, Figure S6). These reciprocal effects are consistent with the floral transition acting as a developmental switch that shapes the balance between vegetative and reproductive growth (Moraes et al., 2019; Shalit et al., 2009). Plants with accelerated flowering completed reproductive development and produced germinable seeds under the conditions tested (Figure S10), indicating that enhanced *FT* input can shorten the time to seed production without abolishing basic reproductive competence. However, the accompanying effects on vegetative growth and final plant stature suggest that quantitative fine-tuning of *FT* activity, rather than constitutive activation, will be important for breeding applications. ALSV-mediated VOX and VIGS, demonstrated here and in previous studies of diverse quinoa traits (Kobayashi and Fujita, 2024; Kobayashi et al., 2025; Ogata et al., 2021), provide an effective platform for such functional testing in this allotetraploid crop while stable transformation and genome editing remain technically challenging.

Several aspects of the present study should be interpreted with caution. Virus-mediated approaches involve transient expression or silencing, and their effects may vary depending on infection efficiency, developmental timing, and target specificity, particularly in highly similar homoeologous gene families. The endogenous role of *CqFT2A*/*2B* remains unresolved because independent silencing could not be achieved. Heterologous expression in *Arabidopsis* may also fail to capture species-specific differences in interacting partners or expression thresholds. Nevertheless, the convergence of quinoa VOX, quinoa VIGS, *Arabidopsis* overexpression, domain-swapping analyses, and cross-germplasm validation supports the conclusion that *CqFT1A* and *CqFT1B-1* are the major florigenic activators in quinoa. More broadly, this study demonstrates that direct functional testing is critical for resolving florigen activity in genetically complex orphan crops and provides a basis for modulating flowering time and generation turnover in climate-resilient quinoa.

## 4. Experimental procedures

### 4.1 Plant Materials and Growth Conditions

Quinoa (*Chenopodium quinoa* Willd.) inbred lines were grown as described previously (Mizuno et al., 2020; Ogata et al., 2021), with minor modifications as described below. All quinoa lines used in this study were inbred lines established or maintained by repeated self-pollination from the indicated accessions or source materials. During seed propagation at JIRCAS, inflorescences were covered with non-woven pollination bags (Rizo, Tsukuba, Japan) to prevent cross-pollination, as described previously (Yasui et al., 2016). The quinoa inbred lines used in this study are listed in Table S1.

For expression analysis of *CqFT* genes in inbred lines, quinoa plants were grown in a growth chamber under a 12-h light/12-h dark photoperiod at 22°C during the light period and 16°C during the dark period, with 60 ± 10% relative humidity and a light intensity of 75 ± 25 µmol photons m^−2^ s^−1^. Most ALSV-mediated VOX and VIGS assays were performed using the lowland-type J082 line in growth chambers (LH-350S/410S; Nippon Medical & Chemical Instruments, Osaka, Japan) under a 16-h light/8-h dark photoperiod at 22°C during the light period and 20°C during the dark period, with 60 ± 10% relative humidity and a light intensity of 150 µmol photons m^−2^ s^−1^. ALSV-mediated assays using late-flowering highland quinoa lines were performed under the same temperature, humidity, and light-intensity conditions, except that plants were grown under a short-day photoperiod of 12-h light/12-h dark photoperiod at 22°C/20°C, respectively) to promote flowering.

*Arabidopsis thaliana* L. accession Columbia-0 (Col-0, CS60000) was used for *Agrobacterium*-mediated transformation. Transgenic *Arabidopsis* lines were generated by the floral dip method (Clough and Bent, 1998) using *Agrobacterium tumefaciens* strain GV3101 (pMP90) carrying the helper plasmid pSoup (Hellens et al., 2000). Seeds of wild-type Col-0 and transgenic lines were surface-sterilized and germinated on germination medium (GM) agar plates containing 1× Murashige and Skoog (MS) salts (Fujifilm Wako, Tokyo, Japan), 1×Gamborg’s vitamin solution (Sigma-Aldrich Japan, Tokyo, Japan), 3% (w/v) sucrose, 0.05% (w/v) MES-KOH (pH 5.7), 0.8% (w/v) agar (Fujita et al., 2005). Plates were stratified at 4°C for 2 days before transfer to 22°C for germination, and the day of transfer was defined as germination day 0. Plants were grown in climate chambers (LH-350S/410S) under controlled conditions with a 16-h light/8-h dark photoperiod at 22°C and a light intensity of 50 ±10 µmol photons m^−2^ s^−1^. For selection and growth of transgenic lines, GM agar medium was supplemented with 25 µg mL^−1^ hygromycin.

### 4.2 Total RNA Isolation and RT-PCR Analysis

Total RNA isolation and complementary DNA (cDNA) synthesis were performed as described previously (Ogata et al., 2017, 2021). Reverse transcription PCR (RT-PCR) was performed using PrimeSTAR GXL DNA polymerase (Takara Bio, Otsu, Shiga, Japan). Primer pairs used for RT-PCR are listed in Table S3. Quantitative RT-PCR (RT-qPCR) was performed to analyze transcript levels, as described previously (Kobayashi et al., 2025; Nagatoshi et al., 2023; Ogata et al., 2021). RT-qPCR was performed using TB Green II (Takara Bio) and a QuantStudio 7 system (Thermo Fisher Scientific, MA, United States), and transcript levels were quantified using the standard curve method. Expression levels were normalized to *CqUBQ10* or 18S *rRNA*, as indicated in the corresponding figure legends. Primer pairs used for RT-qPCR are listed in Table S3. The number of biological replicates is described in each figure legend. For the homoeologous pairs *CqFT1A*/*CqFT1B-1* and *CqFT2A*/*CqFT2B*, transcript levels were detected collectively using primer sets capable of amplifying both homoeologs (Table S3). For expression analysis of *CqFT* genes in the quinoa Kd line, leaves and flower buds were sampled from plants at 1 to 7 weeks after sowing (was). Cotyledons at 1 was, first true leaves at 2–7 was, upper expanded leaves at 3–7 was, and flower buds at the shoot apex at 7 was were collected approximately 3 h after the onset of the light period. Expression levels across different growth stages and organs were normalized to 18S *rRNA*.

### 4.3 Molecular Cloning and Plasmid Construction

To isolate the coding sequences (CDSs) of five *CqFT* genes (*CqFT1A*, *CqFT1B-1*, *CqFT1B-2*, *CqFT2A* and *CqFT2B*) from the standard quinoa inbred line Kd, PCR primers were designed in the 5ʹ and/or 3ʹ untranslated regions (UTRs) based on published CDS and genome sequence information (Jarvis et al., 2017; Yasui et al., 2016). RT-PCR was performed using cDNA prepared from the Kd inbred line. PCR products were cloned into the pGEM-T Easy vector (Promega, Madison, WI, USA), and multiple independent clones were verified by Sanger sequencing. The confirmed sequences were deposited in the DDBJ/EMBL/GenBank databases; accession numbers are listed in section 4.8 and Table S2.

ALSV-RNA2 vectors for VOX and VIGS of *CqFT* genes were constructed using the pEALSR2L5R5 plasmid as previously described (Ogata et al., 2017, 2021). Primer pairs, template DNA, restriction sites used for cloning, and the sizes and sequences of inserts in the ALSV-RNA2 vector are listed in Table S4. The full-length CDS of *AtFT* was amplified from *Arabidopsis* Col-0 cDNA and cloned into pEALSR2L5R5 to generate the *AtFT*-VOX construct. The GFP-VOX vector has been described previously (Ogata et al., 2021).

For domain-swapping analysis, N-terminal or C-terminal variable regions of *CqFT2A* and *CqFT1B-2* were replaced with the corresponding regions of *CqFT1A*. For N-terminal swaps, the N-terminal variable regions of *CqFT2A* and *CqFT1B-2* were replaced with the corresponding 17-amino-acid region of *CqFT1A*, generating *CqFT2A*-N and *CqFT1B2*-N, respectively. For C-terminal swaps, the C-terminal variable regions including segment C in exon 4, which contains the LYN/IYN triplet (Ahn et al., 2006), and the subsequent C-terminal region were replaced with the corresponding 34-amino-acid region of *CqFT1A*, generating *CqFT2A*-C and *CqFT1B2*-C, respectively. The swapped CDSs were synthesized and cloned into the corresponding ALSV-VOX and CaMV 35S overexpression vectors.

To examine whether UTR-derived trigger fragments could improve the specificity of *CqFT1* and *CqFT2* silencing, gene-specific UTR fragments derived from *CqFT1B-1* and *CqFT2B* were used to generate ALSV-based VIGS constructs. Two vector configurations were tested. Single-copy trigger fragments, designated *CqFT1B1*_5′UTR(x1) and *CqFT2B*_3′UTR(x1), were inserted into the cloning site within the 3′ UTR of the ALSV-RNA1 vector pEALSR1-3′SM (Yamagishi et al., 2014). In parallel, multicopy trigger fragments, designated *CqFT1B1*_5′UTR(x3) and *CqFT2B*_3′UTR(x2), were inserted into the ALSV-RNA2 vector pEALSR2L5R5. UTR fragments of *CqFT1B-1* and *CqFT2B* were amplified from the corresponding cDNA clones. For insertion into pEALSR2L5R5, UTR fragments were designed or concatenated so as to maintain the ALSV-RNA2 open reading frame. Details of the UTR-derived VIGS constructs are listed in Table S4.

To generate CaMV 35S promoter-driven overexpression constructs for *Arabidopsis* transformation, pGreen II 0129 (Hellens et al., 2000) -based pGH35S-sGFP vector (Fujita et al., 2009) was modified. The original pGH35S-sGFP vector was used to generate GFP-overexpressing control lines. To construct a multicloning-site vector, an oligonucleotide duplex containing tandem restriction sites (5′-TCTAGAGGATCCATCGATGTCGACCTCGAGGGTACCGCTCCCGGG-3′; XbaI-BamHI-ClaI-SalI-XhoI-KpnI-SmaI) was inserted into the XbaI-SmaI sites between the CaMV 35S promoter and the GFP CDS, generating pGH35S-MCS-sGFP. The GFP CDS was then removed by digestion with XhoI and EcoRV, and PCR-amplified CDSs of *AtFT* or *CqFT* genes, digested with XhoI/EcoRV or XhoI/SmaI, were subcloned into the resulting pGH35S-MCS backbone. The *AtFT* and *CqFT* CDSs were amplified using the primer pairs and ALSV vector clones listed in Table S5. All plasmid constructs listed in Tables S4 and S5 were verified by Sanger sequencing to confirm the inserted sequences.

### 4.4 Virus Inoculation

Quinoa plants were inoculated with ALSV as previously described (Ogata et al., 2017, 2021), with minor modifications. Briefly, purified plasmids containing recombinant pEALSR2L5R5 constructs were mixed with the pEALSR1 plasmid. For constructs based on ALSV-RNA1, recombinant pEALSR1-3′SM plasmids were mixed with the pEALSR2L5R5 plasmid. For the ALSV empty-vector (EV) control, a plasmid mixture of pEALSR1 and empty pEALSR2L5R5 was used. Plasmid DNA mixtures were rub-inoculated onto leaves of the quinoa Iw line using 600-mesh carborundum (Nacalai Tesque, Kyoto, Japan; Ogata et al., 2021). Systemically infected upper leaves showing chlorotic spot symptoms were used to prepare inocula for secondary inoculation of Iw plants. The presence of recombinant ALSV in secondary-inoculated Iw plants was confirmed by symptom development and RT-PCR, as described previously. Systemically infected upper leaves of secondary-inoculated Iw plants were then used to prepare recombinant ALSV inocula for inoculation of other quinoa inbred lines (Ogata et al., 2021). Systemically infected upper leaves from Iw plants inoculated with ALSV extraction buffer (0.1 M Tris-HCl, pH 8.0, 0.1 M NaCl, 5 mM MgCl_2_) were used to prepare mock inoculum. For VOX and VIGS assays, true leaves of the indicated quinoa inbred lines were rub-inoculated with recombinant ALSV inocula prepared from infected Iw source plants using 600-mesh carborundum. Unless otherwise stated, plants were inoculated at 14 days after sowing. Systemically infected upper leaves were sampled at the indicated time points for RT-qPCR analysis to confirm transgene expression or target transcript reduction.

### 4.5 Phenotypic Measurements

For quinoa, bolting time was recorded as the number of days after sowing until visible stem elongation was observed, and flowering time was recorded as the number of days after sowing until at least one flower bud opened. For *Arabidopsis*, bolting time was recorded as the number of days after germination day 0, defined as the day of transfer to 22°C, until visible stem elongation was observed. Flowering time in *Arabidopsis* was recorded as the number of days after germination day 0 until the first flower opened. Total leaf number at flowering was counted at the time of flowering.

For analyses of plant architecture in quinoa, plant height was measured from the soil surface to the shoot apex at the indicated time points. Node number and internode length were measured on the main stem. Seed production and germination capacity of progeny seeds from *AtFT*-VOX-inoculated quinoa plants were evaluated by harvesting mature seeds and monitoring their germination under the growth-chamber conditions described above. Germination was scored at 10 days after sowing. When indicated, the presence of ALSV in progeny seedlings was examined to assess seed transmission.

### 4.6 Sequence, Phylogenetic, and Structural Analyses

Coding sequence and amino acid sequence alignments of FT homologs were performed using ClustalW or Clustal Omega, as indicated in the corresponding figure legends. Phylogenetic trees based on coding sequences or amino acid sequences were constructed in MEGA11 using the neighbor-joining method with 1,000 bootstrap replicates (Tamura et al., 2021). Protein structure predictions for selected FT proteins, including native and domain-swapped CqFT proteins, were performed using ColabFold v1.5.2 with MMseqs2 (Mirdita et al., 2022). The top-ranked predicted models were used for visualization and interpretation. Prediction confidence was evaluated using per-residue predicted local distance difference test (pLDDT) scores, and structural models were visualized using Mol* Viewer (Sehnal et al., 2021).

### 4.7 Statistical Analysis

All statistical analyses were performed using R software version 4.4.0 (R Core Team, 2024). Sample sizes, numbers of independent lines where applicable, and statistical methods used for each experiment are indicated in the corresponding figure legends. For multiple-group comparisons, one-way ANOVA followed by Tukey’s HSD test was used for all-pairwise comparisons, whereas Dunnett’s test was used for comparisons with a control group, as appropriate. When heterogeneity of variance was detected, Welch’s *t*-test with Bonferroni correction or Dunnett’s T3 test was used, as indicated in the figure legends. Data were log-transformed when necessary to stabilize variance or to enable comparisons across several orders of magnitude. Differences were considered statistically significant at *p* < 0.05 unless otherwise stated.

### 4.8 Accession Numbers

The nucleotide sequences of *CqFT1A*, *CqFT1B-1*, *CqFT1B-2*, *CqFT2A*, and *CqFT2B* have been deposited in the DDBJ/EMBL/GenBank databases under the following accession numbers: *CqFT1A* (LC916266), *CqFT1B-1* (LC916267), *CqFT1B-2* (LC916268), *CqFT2A* (LC916269), and *CqFT2B* (LC916270).

## Author Contributions

TO and YF designed the research. TO performed the experiments. TO and YF wrote and reviewed the manuscript. YF supervised the research.

## Acknowledgments

We thank Prof. Emer. N. Yoshikawa (Iwate Univ.) for kindly providing the ALSV-VIGS vectors and quinoa seeds that are the source of the inbred line Iw. We thank the staff of JIRCAS, M. Toyoshima, N. Hisatomi, Y. Masamura, K. Ozawa, K. Shimizu, Y. Shirai, N. Saito, Y. Nonoue, Y. Takiguchi, A. Aoyama, T. Nada, A. Karasawa, Y. Kida, W. Kawakami, and N. Ohmiya for their excellent technical assistance.

## Data Availability Statement

The nucleotide sequences generated in this study have been deposited in the DDBJ/EMBL/GenBank databases under the accession numbers listed in section 4.8. Other data supporting the findings of this study are available within the article and its Supporting Information or from the corresponding author upon reasonable request.

## Supporting information

Additional supporting information can be found online in the Supporting Information section.

---

**Figure S1.**
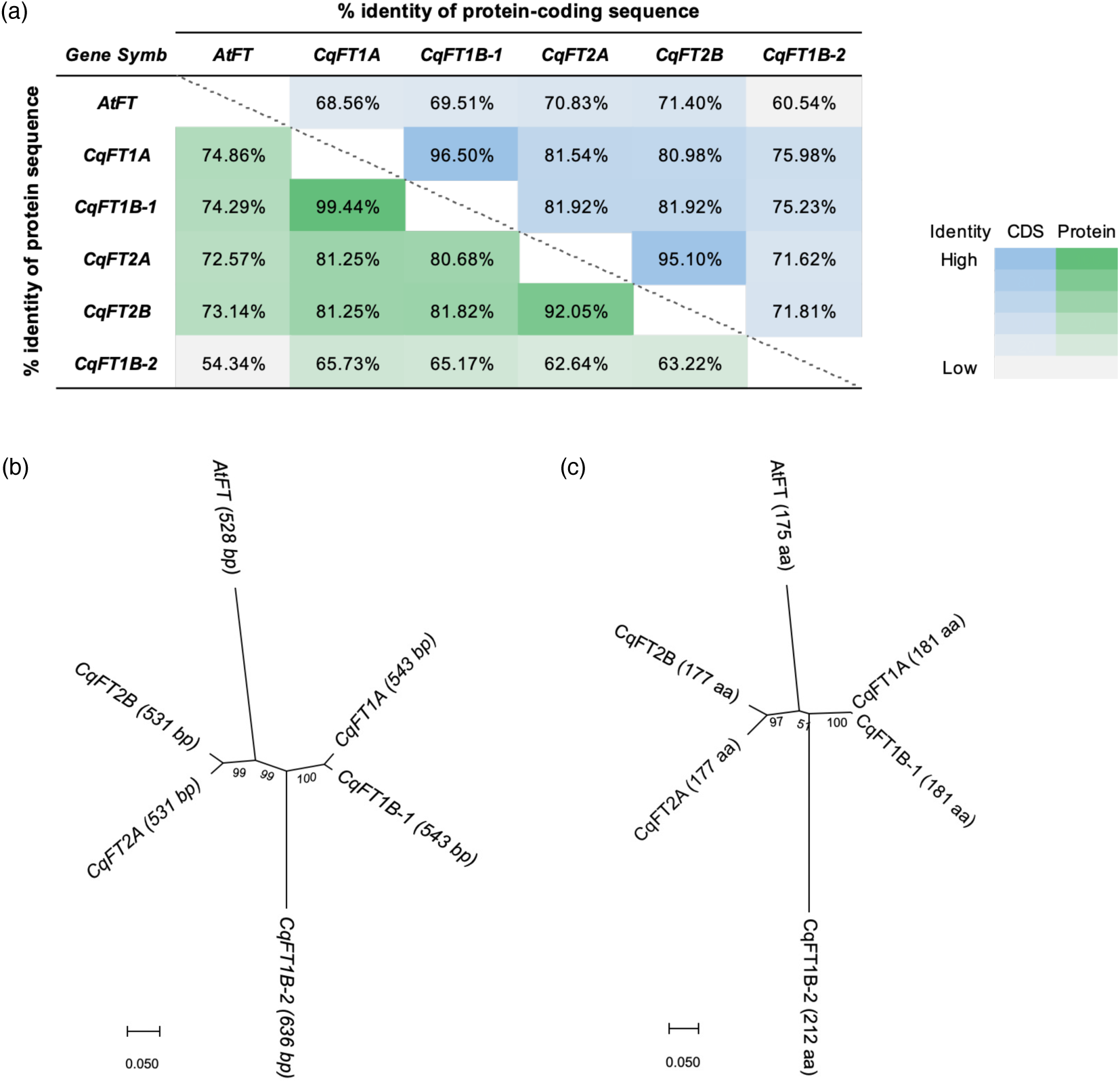
Sequence identity and phylogenetic relationships among quinoa FT homologs. (a) Pairwise identity matrix of protein-coding nucleotide sequences and amino acid sequences among *Arabidopsis thaliana* AtFT and five quinoa CqFT homologs: CqFT1A, CqFT1B-1, CqFT2A, CqFT2B, and CqFT1B-2. Percent identity values are indicated by a color scale. (b, c) Phylogenetic trees based on protein-coding nucleotide sequences (b) and amino acid sequences (c). Sequences were aligned using ClustalW, and phylogenetic trees were constructed using the neighbor-joining method with 1,000 bootstrap replicates in MEGA11.

**Figure S2.**
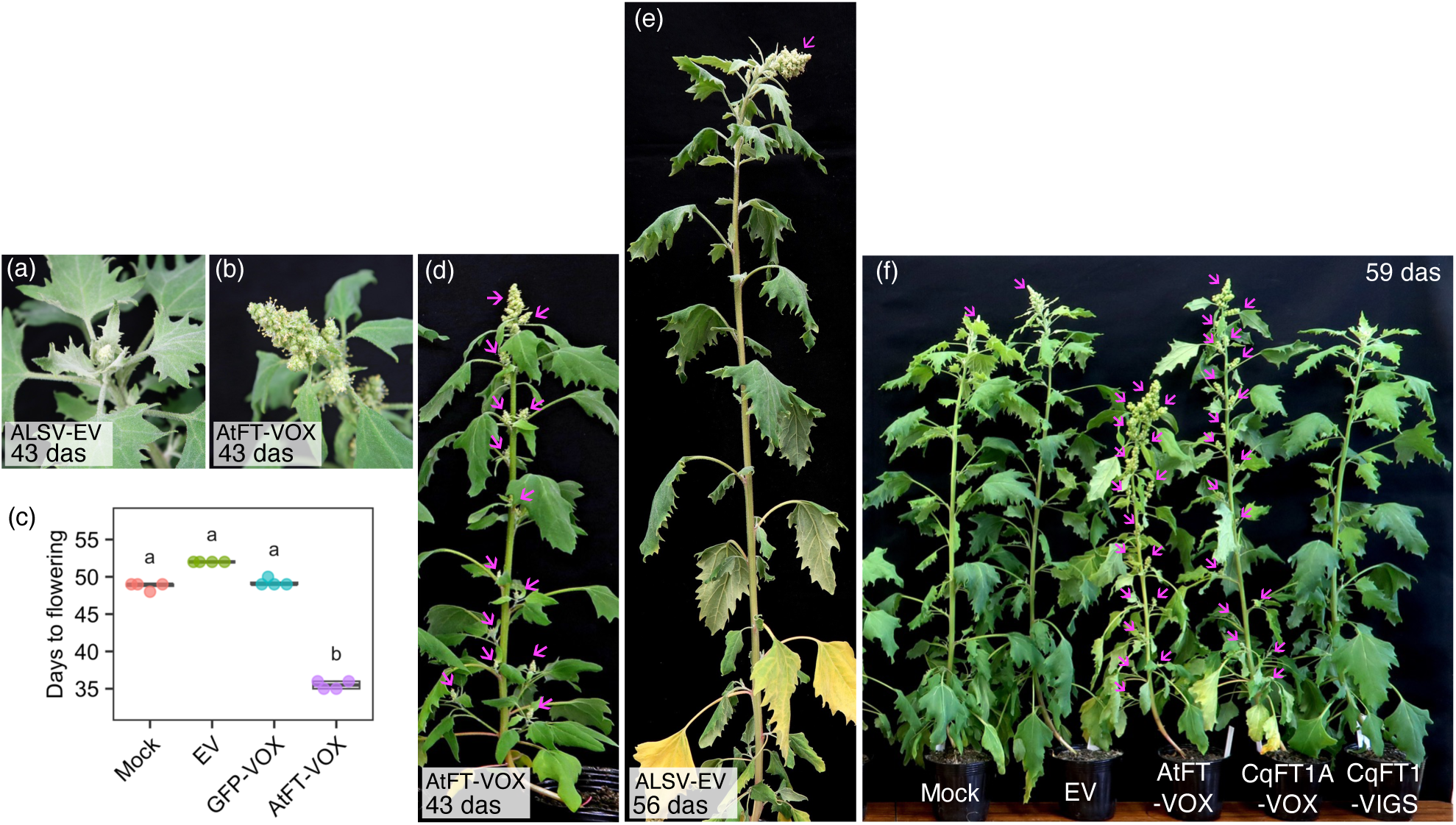
ALSV-mediated overexpression of *Arabidopsis thaliana FT* accelerates flowering in quinoa. (a, b) Representative shoot apices of quinoa plants inoculated with recombinant ALSV inocula corresponding to the empty vector (EV) (a) or *AtFT*-VOX (b) at 43 days after sowing (das). (c) Days to flowering in control and *AtFT*-VOX plants. Days to flowering were recorded as the number of days after sowing until at least one flower bud opened. Buffer-only mock inoculation, EV, and GFP-VOX were used as controls. Data are presented as mean ± SD with individual values (*n* = 4 plants). Different letters indicate significant differences (*p* < 0.001, Welch’s t-test with Bonferroni correction). (d, e) Representative whole plants subjected to *AtFT*-VOX at 43 das (d) or EV at 56 das (e). (f) Representative whole plants subjected to the indicated VOX or VIGS treatments at 59 das. Arrows indicate open flower buds at the time of photography.

**Figure S3.**
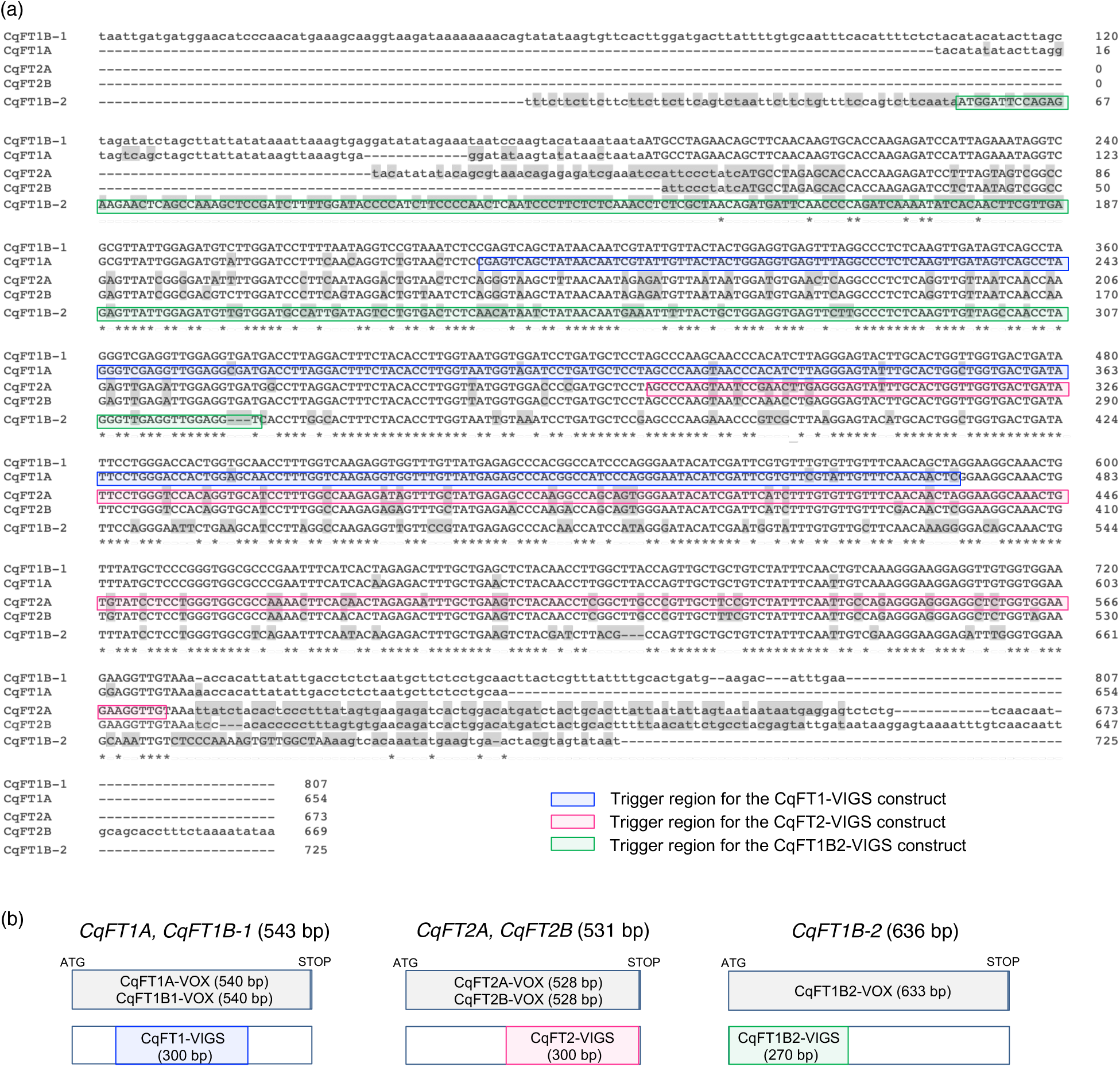
Sequence alignment and trigger map for ALSV vector construction. (a) Sequence alignment of cDNA sequences of quinoa *FT* homologs (*CqFT1A*, *CqFT1B-1*, *CqFT2A*, *CqFT2B*, and *CqFT1B-2*). Primer pairs were designed in untranslated regions (UTRs) based on database sequence information, and RT-PCR was performed using cDNA from the quinoa Kd line. Amplified sequences, with primer-derived regions trimmed, were aligned using Clustal Omega. Coding regions and UTRs are shown in uppercase and lowercase letters, respectively. Identical nucleotides among the five genes are indicated by asterisks (*), and nucleotides differing from *CqFT1B-1* are highlighted in gray. Trigger regions used for *CqFT1*-VIGS, *CqFT2*-VIGS, and *CqFT1B-2*-VIGS are outlined in blue, magenta, and green, respectively. (b) Schematic diagrams of coding sequences and trigger regions cloned into ALSV vectors. For VOX constructs, full-length coding sequences without stop codons were cloned into the pEALSR2L5R5 vector. For VIGS constructs, in-frame coding sequence fragments without stop codons were cloned into the pEALSR2L5R5 vector: 300-bp fragments for *CqFT1*- and *CqFT2*-VIGS and a 270-bp fragment for *CqFT1B-2*-VIGS.

**Figure S4.**
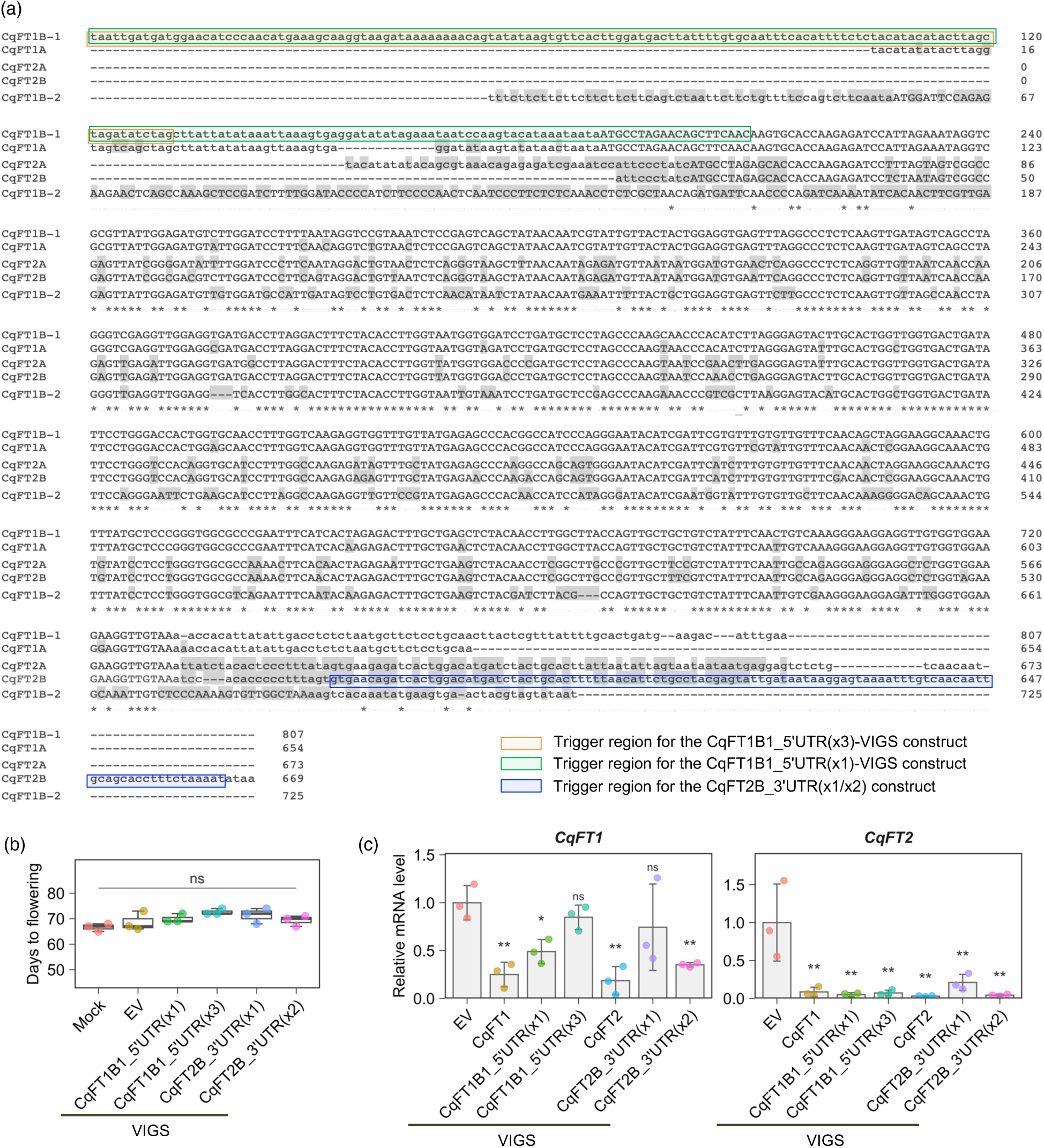
UTR-derived VIGS fragments did not confer effective gene-specific silencing of *CqFT1* or *CqFT2*. (a) Multiple alignment of quinoa *FT* homologs, based on the same alignment shown in Figure S3. UTR-derived trigger regions inserted into ALSV vectors are outlined in color. For CqFT1B1_5′UTR(x1)-VIGS and CqFT2B_3′UTR(x1)-VIGS, single copies of the trigger sequences were inserted into the pEALSR1-3′SM vector. For CqFT1B1_5′UTR(x3)-VIGS and CqFT2B_3′UTR(x2)-VIGS, triple or double copies of the trigger sequences were inserted into the pEALSR2L5R5 vector. (b) Days to flowering in quinoa plants inoculated with recombinant ALSV inocula carrying UTR-derived VIGS trigger fragments. Days to flowering were recorded as the number of days after sowing until at least one flower bud opened. Data are presented as mean ± SD (*n* = 3 plants). ns indicates not significant (*p* > 0.05, one-way ANOVA). (c) RT-qPCR analysis of endogenous *CqFT1* and *CqFT2* transcript levels in quinoa plants inoculated with the indicated recombinant ALSV inocula. Upper leaf samples were collected at 19 days after ALSV inoculation. *CqUBQ10* was used for normalization. Data are presented as mean ± SD (*n* = 3 biological replicates). Statistical significance versus the EV control was determined using Dunnett’s test (\**p* < 0.05, \*\**p* < 0.01; ns, not significant). Buffer-only mock inoculation and EV were used as negative controls, whereas *CqFT1*-VIGS and *CqFT2*-VIGS were included as reference VIGS treatments. For *CqFT1* and *CqFT2*, primer pairs detected the combined transcript levels of *CqFT1A*/*CqFT1B-1* and *CqFT2A*/*CqFT2B*, respectively.

**Figure S5.**
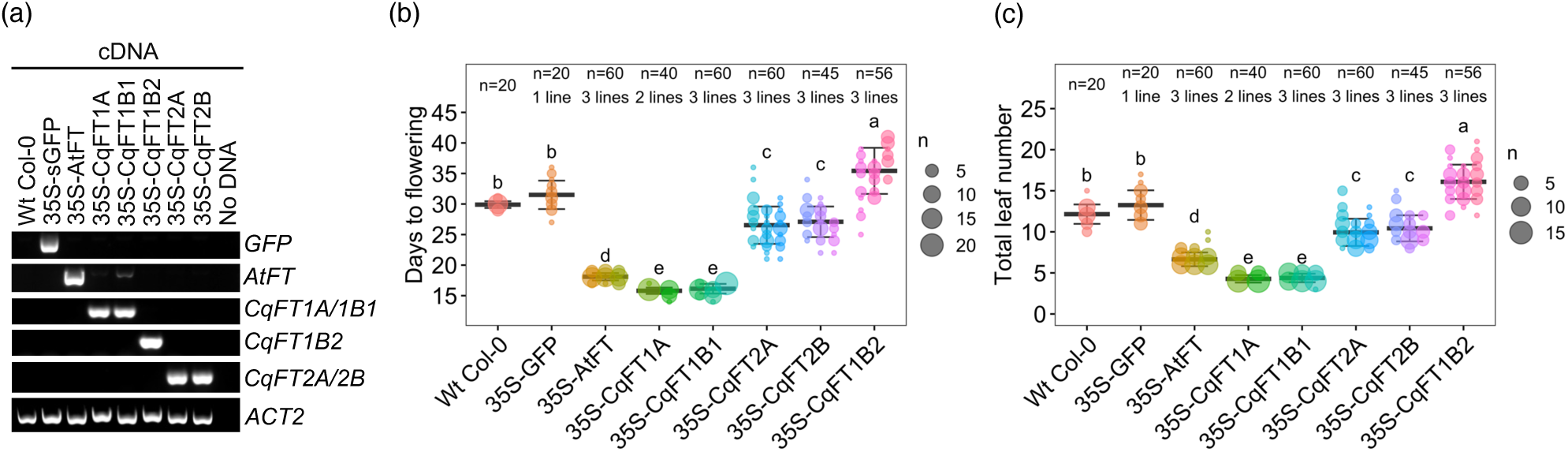
Ectopic overexpression of quinoa *FT* homologs alters flowering time in *Arabidopsis*. *Arabidopsis thaliana* Col-0 was transformed with CaMV 35S promoter-driven *FT* genes. Wild-type (Wt) Col-0 and T3 homozygous GFP-overexpressing lines (35S-GFP) were grown as controls. (a) RT-PCR validation of transgene expression in transgenic *Arabidopsis* lines. *AtACT2* was amplified as an internal control. (b, c) Days to flowering from germination day 0 (b) and total leaf number at flowering (c) were measured for each line, in addition to days to bolting shown in Figure 3. Two or three independent T2 lines were analyzed per construct, except for Wt Col-0 and 35S-GFP controls. Data are presented as mean ± SD of all plants analyzed, and individual lines are shown in different colors. The number of lines and total plants (*n*) are indicated at the top. Different letters indicate significant differences (*p* < 0.001, Dunnett’s T3 test).

**Figure S6.**
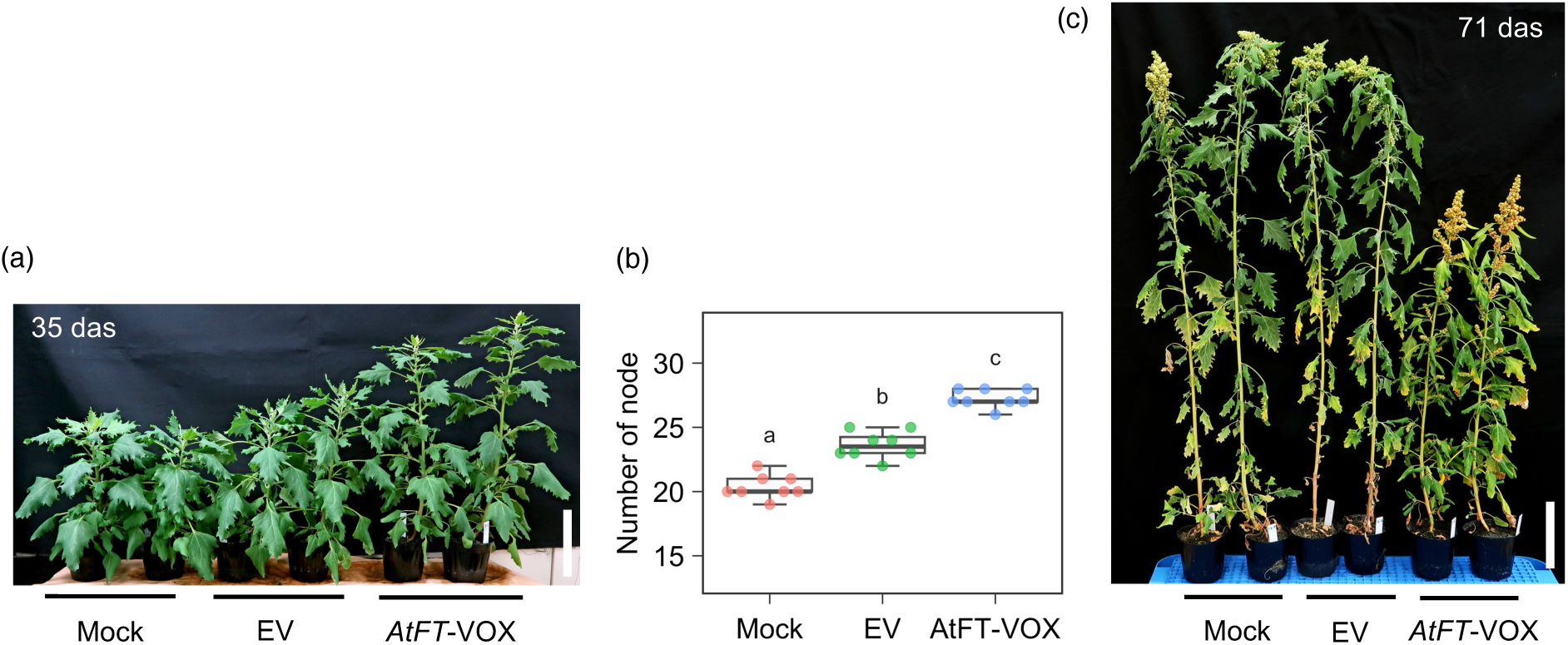
Flowering acceleration by *AtFT*-VOX alters plant height in quinoa. (a, b) *AtFT*-VOX accelerated flowering and increased node number at the early flowering stage. Quinoa J082 plants were subjected to buffer-only mock inoculation, EV, or *AtFT*-VOX. (a) Representative photograph taken at 35 days after sowing (das), when *AtFT*-VOX plants started flowering. (b) Node number at 35 das. Data are presented as mean ± SD with individual values (*n* = 8 plants). Different letters indicate significant differences (*p* < 0.01, Tukey’s HSD test). (c) Representative photograph showing reduced final plant height in *AtFT*-VOX plants. The photograph was taken at 71 das, when EV plants had completed flowering. Scale bars in (a) and (c) represent 10 cm.

**Figure S7.**
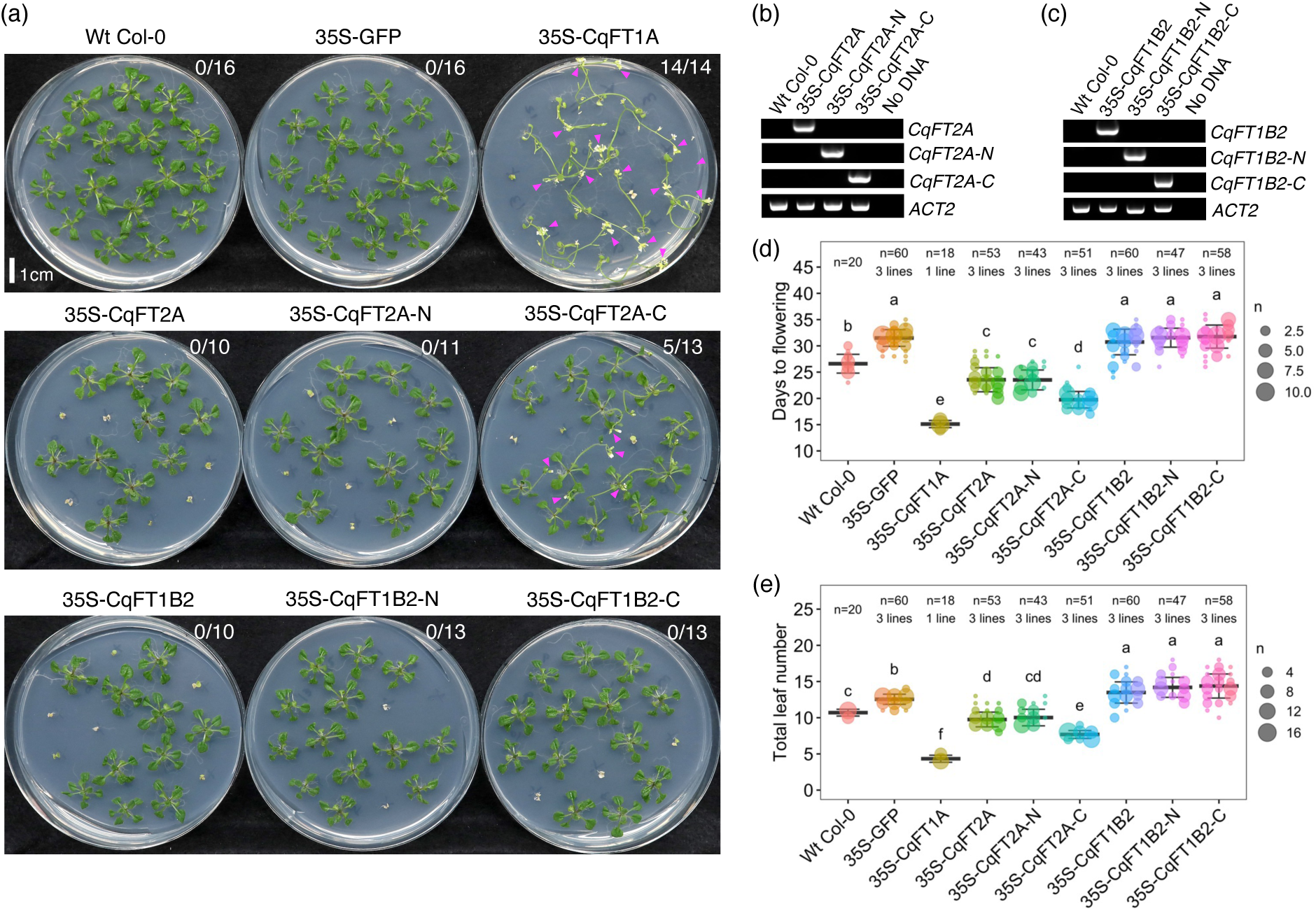
Terminal-region substitutions in CqFT proteins differentially affect flowering-promoting activity in *Arabidopsis*. (a) Representative images of transgenic *Arabidopsis* lines expressing native or N-terminal/C-terminal-swapped forms of *CqFT2A* and *CqFT1B-2* under the control of the CaMV 35S promoter. Photographs were taken 20 days after germination day 0, defined as the day of transfer to 22°C. One representative independent T2 line is shown for each construct. T2 plants were grown on hygromycin-containing medium to exclude null segregants. Wt Col-0 and T3 homozygous 35S-GFP lines were grown as controls. Magenta arrowheads indicate plants flowering at this time point, and the number of flowering plants relative to hygromycin-resistant plants is shown. (b, c) RT-PCR validation of transgene expression in transgenic *Arabidopsis* lines expressing native or domain-swapped *CqFT2A* (b) and *CqFT1B-2* (c) constructs. *AtACT2* was amplified as an internal control. (d, e) Days to flowering from germination day 0 (d) and total leaf number at flowering (e) were recorded for each line, in addition to days to bolting shown in Figure 5g. Three independent T2 lines were analyzed per construct, except for the 35S-*CqFT1A* control line. Data are presented as mean ± SD of all plants analyzed, with individual lines shown in different colors. The number of lines and total plants (*n*) are indicated at the top. Different letters indicate significant differences (*p* < 0.001, Dunnett’s T3 test).

**Figure S8.**
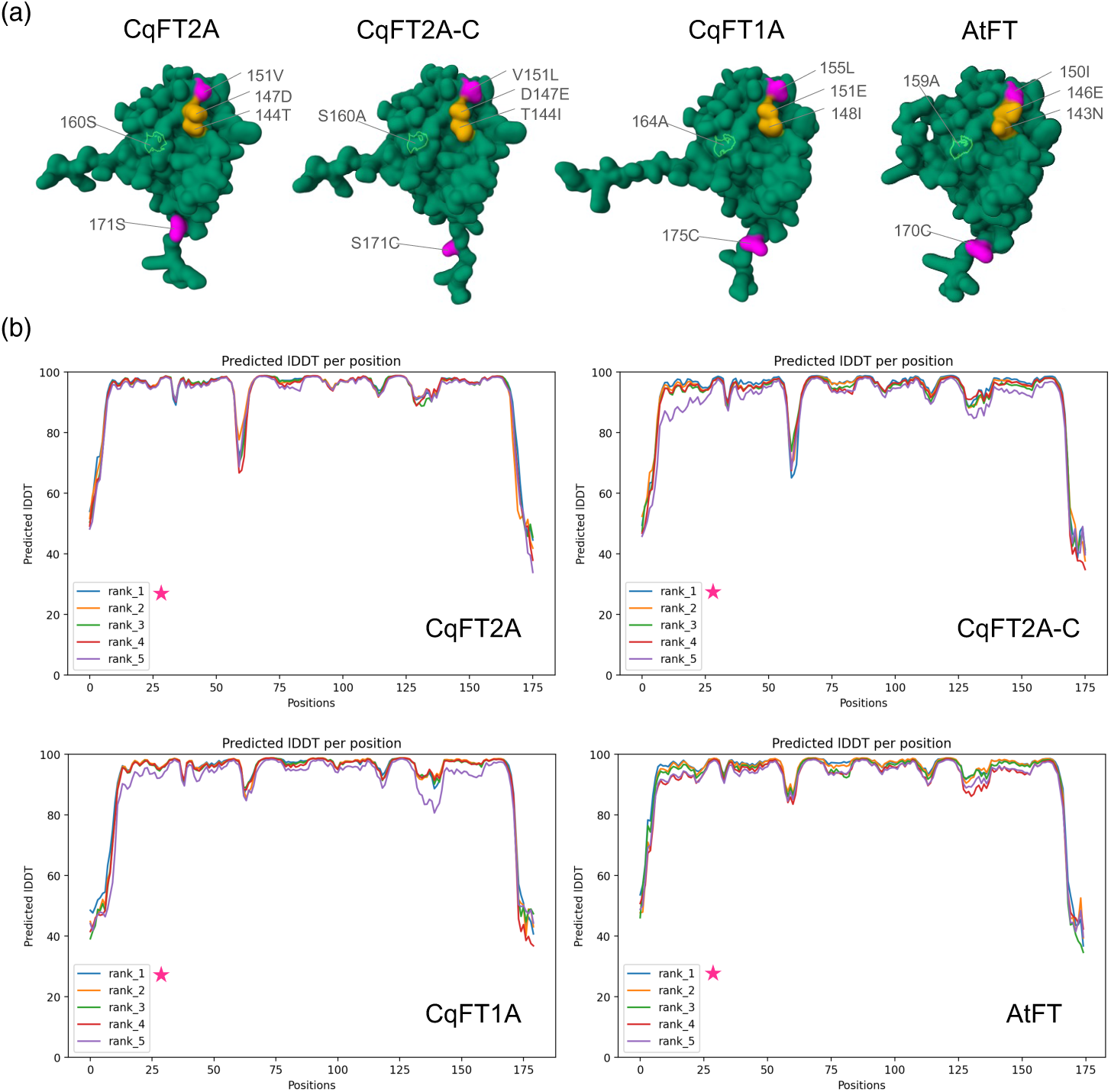
Comparison of ColabFold-predicted protein 3D structures for FT proteins. (a) Predicted three-dimensional structures of CqFT2A, the C-terminal-swapped CqFT2A-C protein, CqFT1A, and AtFT. Protein structures were predicted using ColabFold v1.5.2 with MMseqs2, and the top-ranked predicted models were visualized using Mol* Viewer. Amino acid positions that differ between CqFT2A and CqFT1A within the C-terminal swapping region are indicated: positions 151 and 171 are shown in magenta, positions 144 and 147 in orange, and position 160 in green. (b) Per-residue predicted local distance difference test (pLDDT) plots for the predicted models shown in (a).

**Figure S9.**
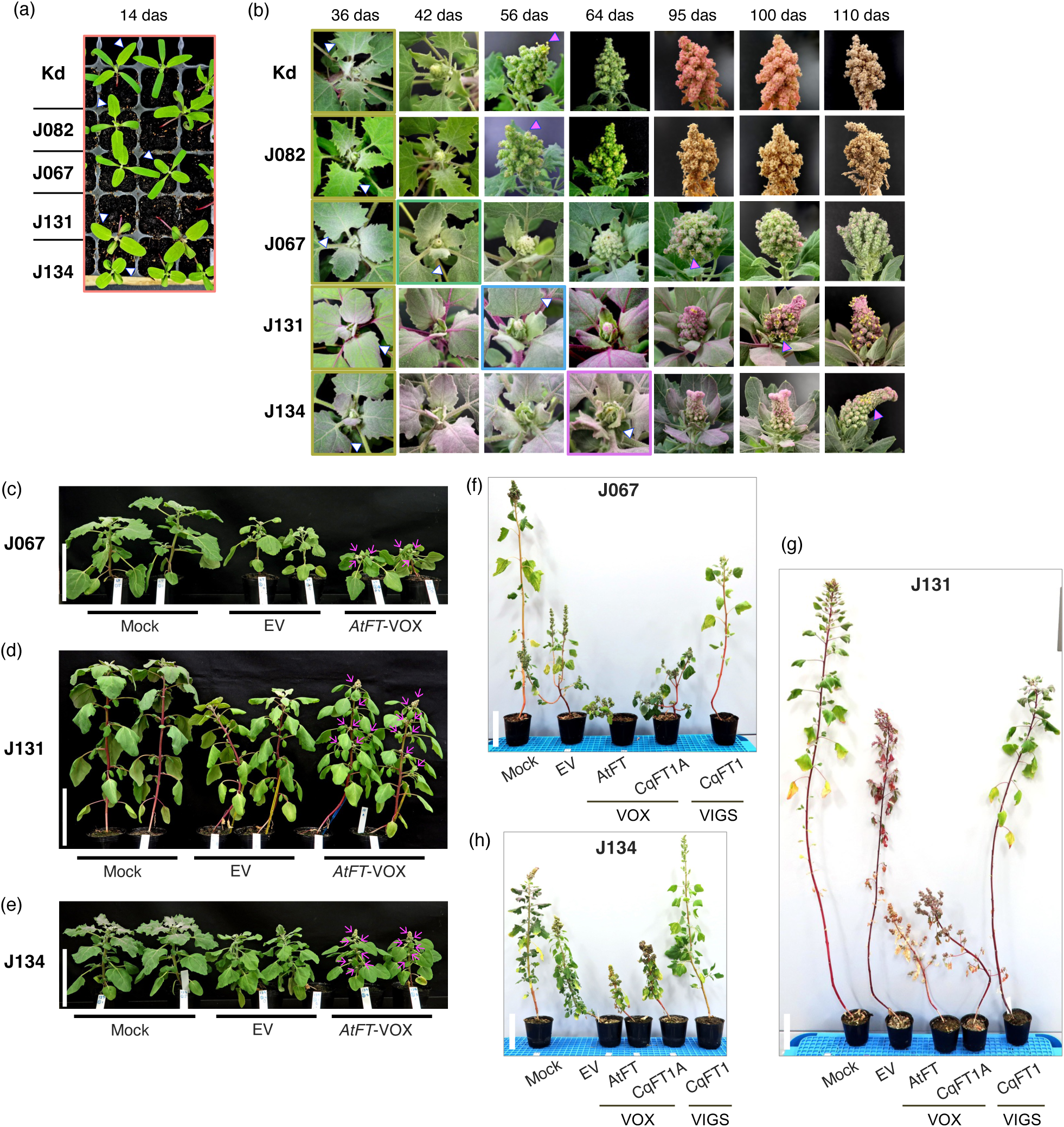
Regulation of flowering by ALSV-mediated modulation of *FT* expression in five quinoa lines. (a, b) Representative photographs of quinoa lines Kd, J082, J067, J131, and J134 at seedling, bolting, and flowering stages. Kd and J082 are lowland lines, whereas J067, J131, and J134 are highland lines. (a) Young seedlings at 14 days after sowing (das). (b) Morphological changes of shoot apices from 36 to 110 das. Plants were grown under a 12-h light period at 22°C and a 12-h dark period at 16°C, with 50% ± 10% relative humidity, and were used for RT-qPCR analysis. Colored lines outline plants at sampling time points. White arrowheads indicate sampled upper true leaves, and magenta arrowheads indicate open flowers. Photographs are not to scale. (c–h) Plant morphology of three late-flowering highland lines subjected to ALSV-mediated VOX or VIGS treatments: J067 (c, f), J131 (d, g), and J134 (e, h). Photographs were taken at 50 das (c–e) or 103 das (f–h). Arrows in (c–e) indicate open flower buds at the time of photography. In J131, plants subjected to *CqFT1A*-VOX and *CqFT1*-VIGS initiated flowering at approximately 50 das and 103 das, respectively. Buffer-only mock inoculation and EV were used as controls. Plants were grown under a 12-h light period at 22°C and a 12-h dark period at 20°C. White scale bars represent 10 cm.

**Figure S10.**
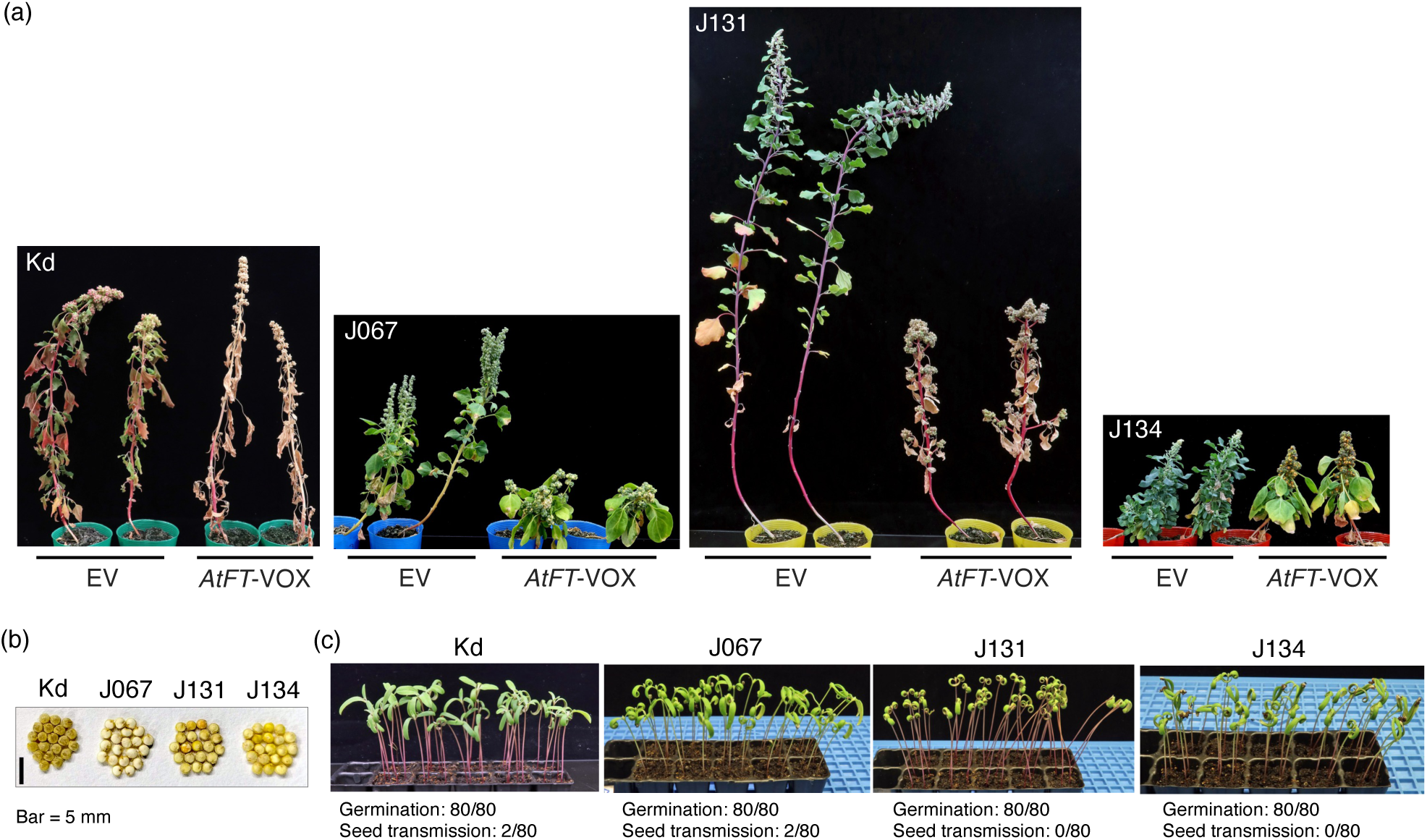
*AtFT*-VOX plants produce viable progeny seeds within a shortened timeframe. (a) Representative photographs of quinoa plants subjected to EV or *AtFT*-VOX at harvest. Photographs were taken at 15 weeks after sowing for Kd and at 17 weeks after sowing for J067, J131, and J134. (b) Mature seeds harvested from *AtFT*-VOX plants of the indicated quinoa lines. Scale bar = 5 mm. (c) Germination assay of progeny seeds harvested from *AtFT*-VOX plants. Photographs were taken at 10 days after sowing. The numbers of germinated seedlings and ALSV-positive seedlings per sown seed are indicated at the bottom.

**Table S1.**
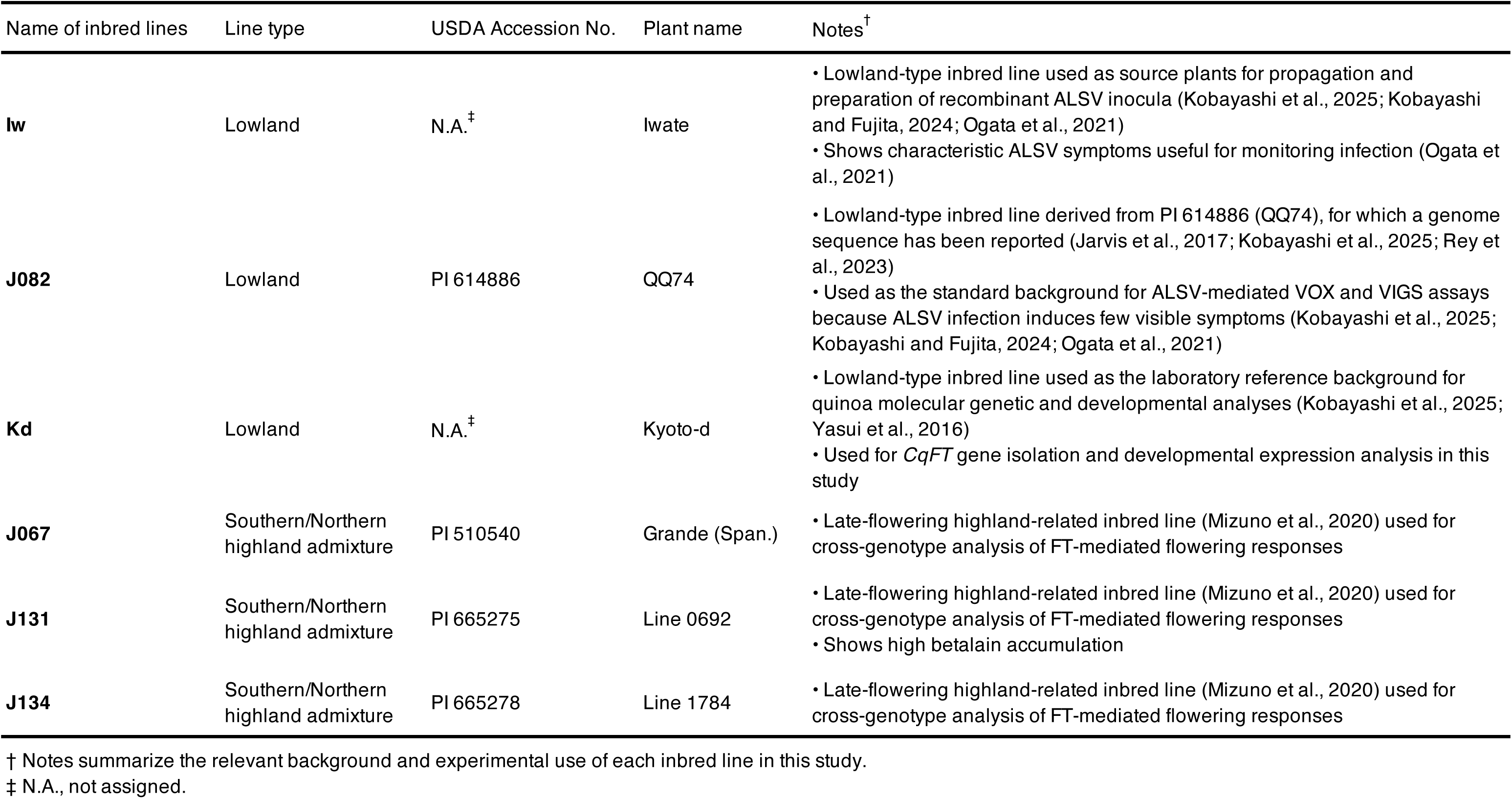
Quinoa inbred lines used in this study.

**Table S2.**
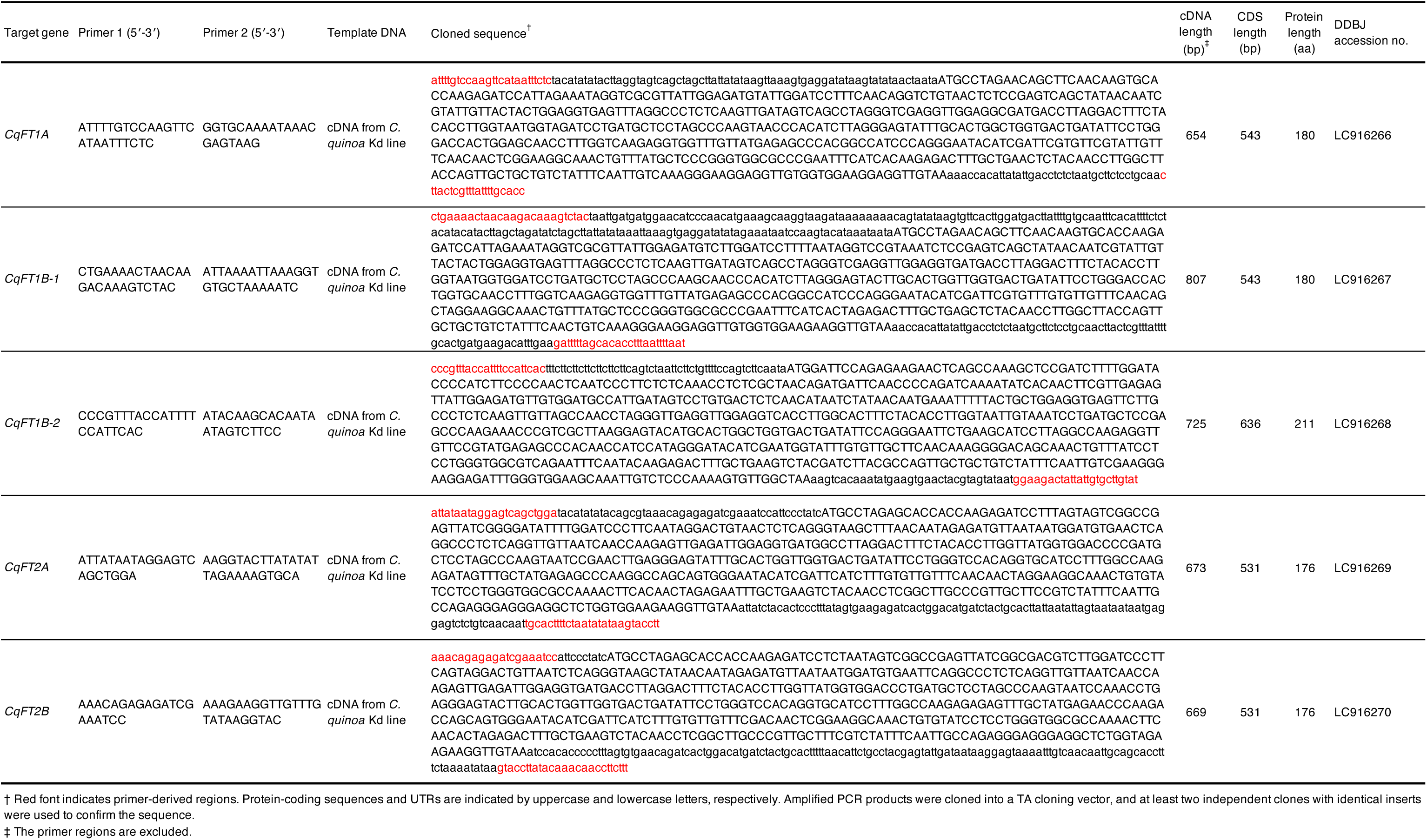
Cloned *CqFT* sequences, accession numbers, and primers used for gene cloning.

**Table S3.**
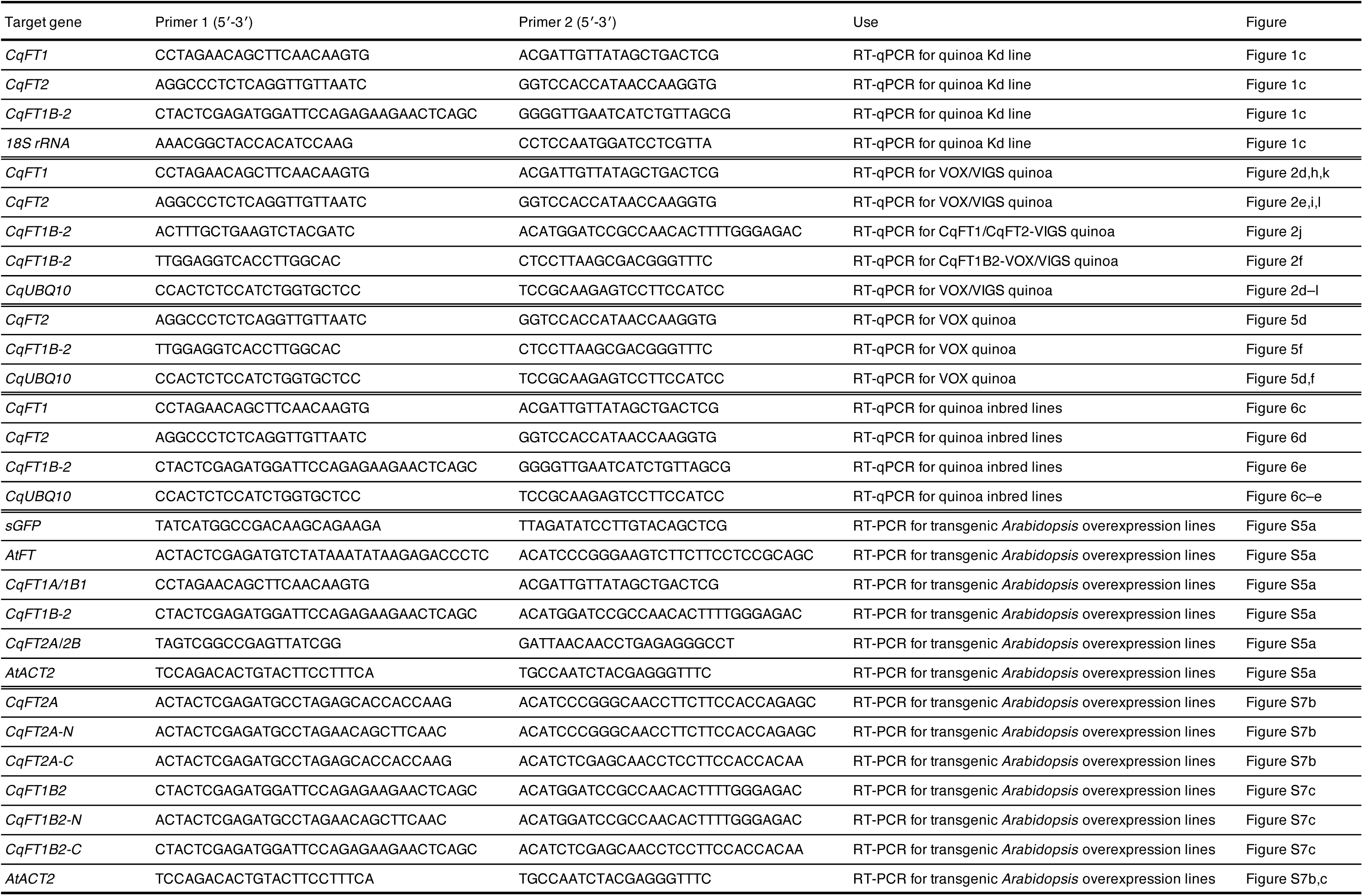
Primers used for RT-PCR and RT-qPCR expression analyses.

**Table S4.**
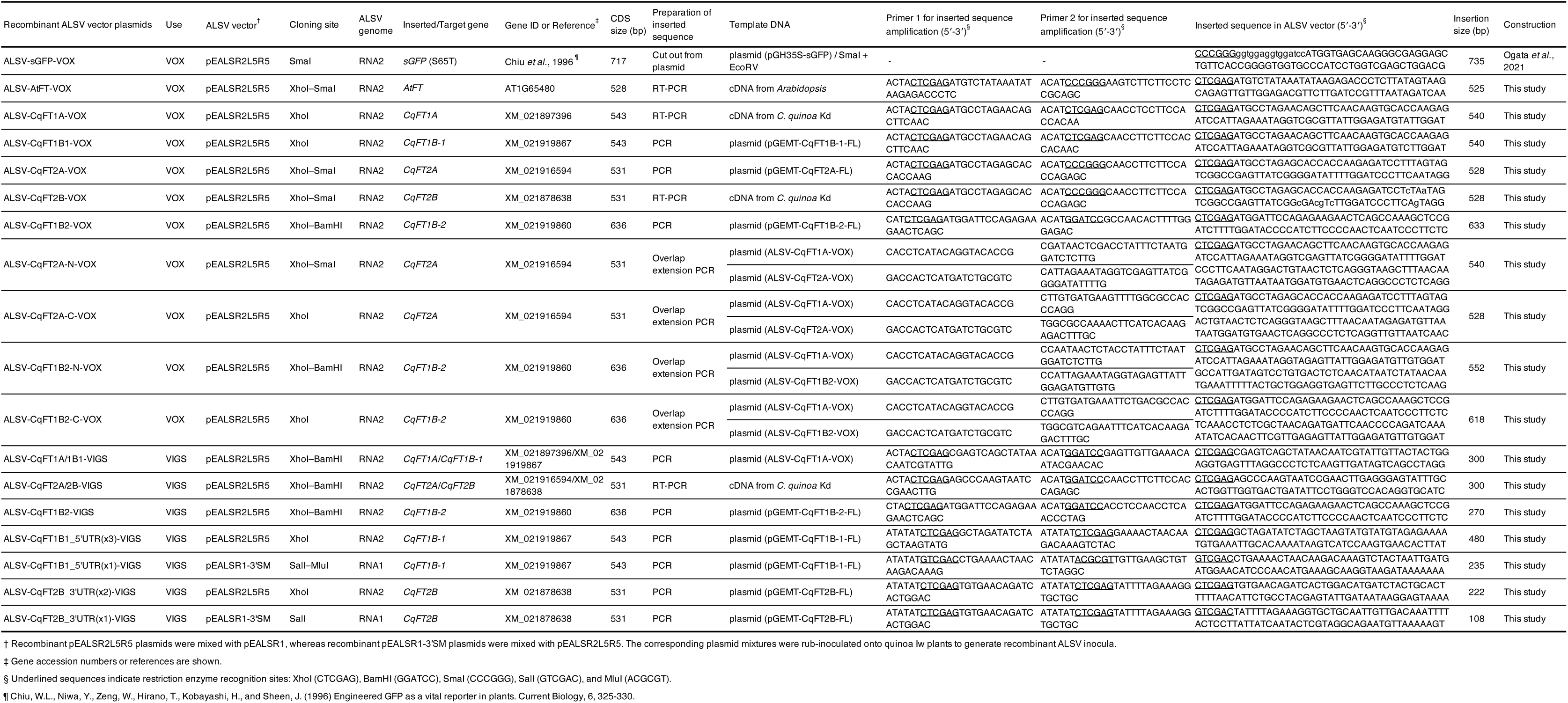
Construction of recombinant ALSV vectors used for VOX and VIGS.

**Table S5.**
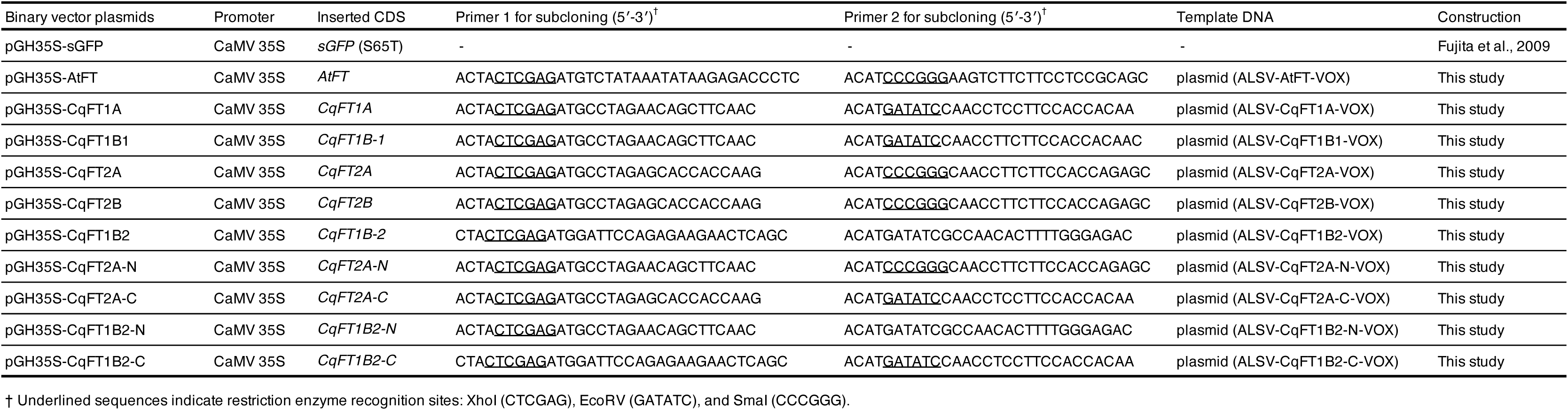
Construction of CaMV 35S promoter-driven plasmids used for *Arabidopsis* transformation.

